# Poly(dA:dT) tracts differentially modulate nucleosome remodeling activity of RSC and ISW1a complexes, exerting tract orientation-dependent and -independent effects

**DOI:** 10.1101/2023.07.25.550574

**Authors:** Roberto Amigo, Fernanda Raiqueo, Estefanía Tarifeño, Carlos Farkas, José L. Gutiérrez

## Abstract

The establishment and maintenance of nucleosome-free regions (NFRs) are prominent processes within chromatin dynamics. Transcription factors, ATP-dependent chromatin remodeling complexes (CRCs) and DNA sequences are the main factors involved. In *Saccharomyces cerevisiae*, CRCs such as RSC contribute to chromatin opening at NFRs, while other complexes, including ISW1a, contribute to NFR shrinking. Regarding DNA sequences, growing evidence point to poly(dA:dT) tracts as playing a direct role in the active processes involved in nucleosome positioning dynamics. Intriguingly, poly(dA:dT) tract-containing NFRs span asymmetrically relative to the location of the tract, by a currently unknown mechanism. In order to get insight into the role of poly(dA:dT) tracts in nucleosome remodeling, we performed a systematic analysis of their influence on the activity of ISW1a and RSC complexes, with focus on their effect when located on linker DNA. Our findings revealed that poly(dA:dT) tracts differentially affect the activity of these CRCs. Moreover, we found differences between the effects exerted by the two alternative tract orientations. Our results support a model for the asymmetric chromatin opening that take place around these sequences.

## Introduction

Regulation of gene expression at the level of transcription requires the coordinated contribution of a large number of factors. Among these are ATP-dependent chromatin remodeling complexes (CRCs), which alter the chromatin landscape at defined genomic regions by evicting histone octamers, altering nucleosome positioning and/or altering their composition (Clapier, 2021). In this context, the position of nucleosomes at gene regulatory regions plays a key role in transcriptional regulation. Genome-wide analyses of nucleosome positioning have uncovered a common chromatin landscape, consisting in a well-positioned nucleosome, called +1 nucleosome, covering the transcription start site (TSS) or few base pairs downstream this site, followed by a nucleosome array that covers part of the gene body. A long stretch of DNA of low nucleosome occupancy is commonly observed upstream the +1 nucleosome, called nucleosome-depleted region (NDR) or nucleosome-free region (NFR). These terms have been commonly used as equivalent terms and we do so in the current report, although it has to be noted that differences between them have been described (Lai & Pugh, 2017). The typical chromatin landscape in the boundaries of TSSs is completed by an also well-positioned nucleosome located upstream the NFR, called the-1 nucleosome (Lai & Pugh, 2017; Clapier, 2021).

It is currently conceived that the players involved in the dynamics of nucleosome positioning are transcription factors, CRCs and DNA sequences (Lai & Pugh, 2017; Brahma & Henikoff, 2020; Barnes & Korber, 2021; Clapier, 2021). Within this context, it has been observed that defined CRCs play opposite roles at gene promoter NFRs. Particularly in the yeast *Saccharomyces cerevisiae*, the RSC complex plays a role in establishment and maintenance of the NFR, while the ISW1a complex performs NFR shrinking (Parnell *et al*, 2015). In addition, other CRCs have been pointed as relevant players in NFR maintenance and positioning of the +1 nucleosome in this species, such as Ino80 and ISW2 (Kubik *et al*, 2019; Oberbeckmann *et al*, 2021b). Regarding the involvement of DNA sequences on nucleosome positioning, early genome-wide analyses assigned a mayor role to them. It was then conceived that this role relies essentially on variations in flexibility of different DNA sequences, which drive histone octamers to preferential translational locations (Segal *et al*, 2006; Kaplan *et al*, 2009). Further studies using *in vitro* reconstitution of the whole *S. cerevisiae* genome into nucleosomes, determined that the obtaining of an *in vivo*-like nucleosome positioning pattern depends essentially on active processes. Thereafter, these studies pointed to transcription factors and CRCs, notably RSC, as the main players involved in the processes determining nucleosome positioning (Wippo *et al*, 2011; Zhang *et al*, 2011; Korber, 2012; Struhl & Segal, 2013). A more marginal and passive role was then assigned to DNA sequences in nucleosome positioning. This passive role of DNA sequences would include mechanical (intrinsic) properties, which determine variations in flexibility of different sequences, and the presence of binding sites for transcription factors that participate in nucleosome remodeling processes (Korber, 2012; Struhl & Segal, 2013). Afterwards, however, a growing number of studies have pointed to DNA sequences as relevant players in the active processes involved in the dynamics of nucleosome positioning. Until now, the participation of DNA sequences in these active processes has been essentially postulated with base on the observed effects of poly(dA:dT) tracts on ATP-dependent nucleosome remodeling activity (Lorch *et al*, 2014; Krietenstein *et al*, 2016; Winger & Bowman, 2017; Barnes & Korber, 2021).

The presence of poly(dA:dT) tracts at gene regulatory regions in *S. cerevisiae* was reported few years after the onset of DNA sequencing. These studies and further analyses demonstrated the involvement of these sequences mainly in transcriptional regulation of constitutive genes, but also in inducible transcription (Struhl, 1985; Iyer & Struhl, 1995). Afterwards, an abundant number of *in vivo* and *in vitro* genome-wide studies have shown the involvement of these tracts in NFR formation and maintenance (Barnes & Korber, 2021; Clapier, 2021). It has been determined that these sequences enhance nucleosome disassembly mediated by the RSC complex (Lorch *et al*, 2014) and influence the nucleosome remodeling activity of CHD1 (Winger & Bowman, 2017). Intriguingly, it has been observed that NFRs harboring a single poly(dA:dT) tract span a region that is asymmetrically distributed around the tract, with a longer stretch of the NFR located 5’ of the polyA sequence (de Boer & Hughes, 2014). In the same context, there is a higher frequency of polyT in the sense strand upstream the midpoint of gene promoter NFRs, while there is enrichment of polyA in the sense strand downstream this midpoint, with the presence of both patterns at individual gene promoters being less frequent than expected (Wu & Li, 2010). Although it is known that RSC has the ability to asymmetrically clear chromatin around poly(dA:dT) tracts (Krietenstein *et al*, 2016), the means by which this biochemical outcome is obtained are currently unknown. In addition, there are no studies comparing the effects of the two alternative orientations of poly(dA:dT) tracts on ATP-dependent nucleosome remodeling activity.

In order to get insight into the role played by poly(dA:dT) tracts in active processes involved in nucleosome positioning, we performed a systematic analysis of the effect of these sequences on ATP-dependent nucleosome remodeling activity. Considering their stimulatory effect on RSC remodeling activity, we aimed to get further insight into the effects of these tracts on the activity of this complex, specifically RSC2 (Schlichter *et al*, 2020). In addition, taking into account the opposite roles played by ISW1a and RSC at NFRs, we also aimed to determine whether these tracts affect the activity of ISW1a and, if so, whether they differentially affect the activity of RSC and ISW1a complexes. To do this, we used a large variety of nucleosome probes harboring these tracts in different orientations and positions relative to the nucleosome core, in addition to probes harboring different lengths of the tracts. Our analyses were mainly focused on the effect displayed by these tracts when located on linker DNA, considering the following criteria: Firstly, poly(dA:dT) tracts of the *S. cerevisiae* genome mainly locate at NFR or linker regions not only *in vivo* but also upon *in vitro* reconstitution of this genome into nucleosomes (Kaplan *et al*, 2009; Krietenstein *et al*, 2016). Secondly, interaction with linker DNA has been described for both complexes, including mechanistic implications of these interactions in the remodeling process. For ISW1a, as for other complexes of the ISWI subfamily, interaction with linker DNA stimulate its remodeling activity and makes this linker the entry DNA in the sliding process (Gangaraju & Bartholomew, 2007; Clapier, 2021; Reyes *et al*, 2021). For the RSC complex, recent structural analyses have characterized a linker DNA-interaction module (DIM) in the complex, with this linker DNA being the exit DNA during nucleosome remodeling (Ye *et al*, 2019; Wagner *et al*, 2020). Our findings revealed that poly(dA:dT) tracts differentially affect the activity of ISW1a and RSC complexes. Moreover, we found that the two alternative orientations of poly(dA:dT) tracts exert differential effects on nucleosome remodeling activity. In the case of RSC, this differential effect implies different outcomes of RSC activity. In addition, we found that poly(dA:dT) tracts orient RSC to select the tract-containing linker as the exit DNA during remodeling. These new properties of poly(dA:dT) tracts might conform the underlying mechanism of the asymmetric chromatin opening that take place around these sequences.

## Results

### Poly(dA:dT) tracts exert a differential effect on the activity of ISW1a and RSC complexes

In this study we used a large number of nucleosome probes that harbor different designs, mainly in terms of location and orientation of poly(dA:dT) tracts (see Table S1 for details). To clearly state the features of every probe, including location and orientation of tracts, we use a set of nomenclature guidelines throughout the whole text. One strand of the Widom’s 601 pronucleosomal sequence is arbitrarily defined as “upper strand” (Fig S1); in some sections, particularly in the Discussion, the upper strand is considered equivalent to the sense strand of genes. The location of a poly(dA:dT) tract is defined as upstream or downstream the nucleosome core (or dyad), relative to the upper strand, when referring to a tract as a whole double-strand entity and also when referring to only one of the tract sequences (polyA or polyT). The orientation of the poly(dA:dT) tract is defined by indicating the presence of polyA (or polyT) in the upper or the lower strand. Tract orientation is also defined in terms of the location of the nucleosome core or dyad relative to 5’ or 3’ end of the polyA sequence (see a summary of these denominations in Fig S1). For easiness, extra-core DNA (aka extranucleosomal DNA) is referred to as “linker DNA” or “linker” throughout the text. Nucleosome probes are called including their main features in the names, such as: location of nucleosome core; presence and length of linker DNA; and presence, length and orientation of poly(dA:dT) tracts. These elements are named relative to the 601 upper strand, following its 5’ to 3’ direction (i.e., from left to right). The <u>n</u>ucleosome <u>c</u>ore is designated as “NC”.

In order to ascertain whether poly(dA:dT) tracts differentially affect the activity of RSC and ISW1a complexes, together with testing the two alternative orientations of these tracts, we first performed remodeling assays comparing a set of three probes reconstituted into mononucleosomes. These 217 bp probes harbor the 147 bp positioning region of the 601 sequence (or the first 145 bp of this sequence), to position the histone octamer upon nucleosome reconstitution, plus 70 (or 72) bp downstream this sequence. These 70 bp become linker DNA after reconstitution (0-NC-70 probes, Fig 1A, left panel). Two of these probes contain a 15 bp poly(dA:dT) tract located immediately downstream the 145 bp 601 sequence; one of them harbors the polyA sequence in the upper strand and the other one harbors this sequence in the lower strand. Under this design, most of the tract becomes part of linker DNA upon nucleosome reconstitution (Fig 1A). The sliding activity of these complexes was visualized by electrophoresis in non-denaturing gels, followed by quantification of the slid over total mononucleosome signal ratio (Fig 1B). In the case of RSC, its sliding activity involves generation of a fast migrating nucleosomal band that has been characterized as entailing mobilization of the histone octamer up to 51 bp beyond a DNA end (Flaus & Owen-Hughes, 2003). Interestingly, the presence of these tracts hindered the sliding activity of ISW1a (Fig 1C, compare lane 2 to 4 and 6, and the corresponding probes in the graph). Moreover, we found that there is a difference between the extents of inhibition given by the orientation of the homopolymeric tract, with the strongest inhibition corresponding to the presence of polyA in the lower strand (Fig 1C). Conversely, in the case of RSC-as expected from previous studies (Lorch *et al*, 2014)-the presence of the tracts enhanced its sliding activity (Fig 1D, compare lane 4 to 6 and 8, and the corresponding probes in the graph). Here, we also found a differential effect given by the orientation of the tract, with the strongest stimulation observed in the probe harboring polyA in the lower strand (Fig 1D). This difference in the stimulatory effect on RSC activity only apply to its sliding activity, as determined by the ratio of slid over total mononucleosome signal, and not to other biochemical outcomes that are addressed below. Thus, the orientation that exerted the strongest stimulatory effect on the sliding activity of RSC was the same that resulted in the strongest inhibitory effect on the sliding activity of ISW1a (Figs 1C and 1D, respectively).

**Figure 1.**
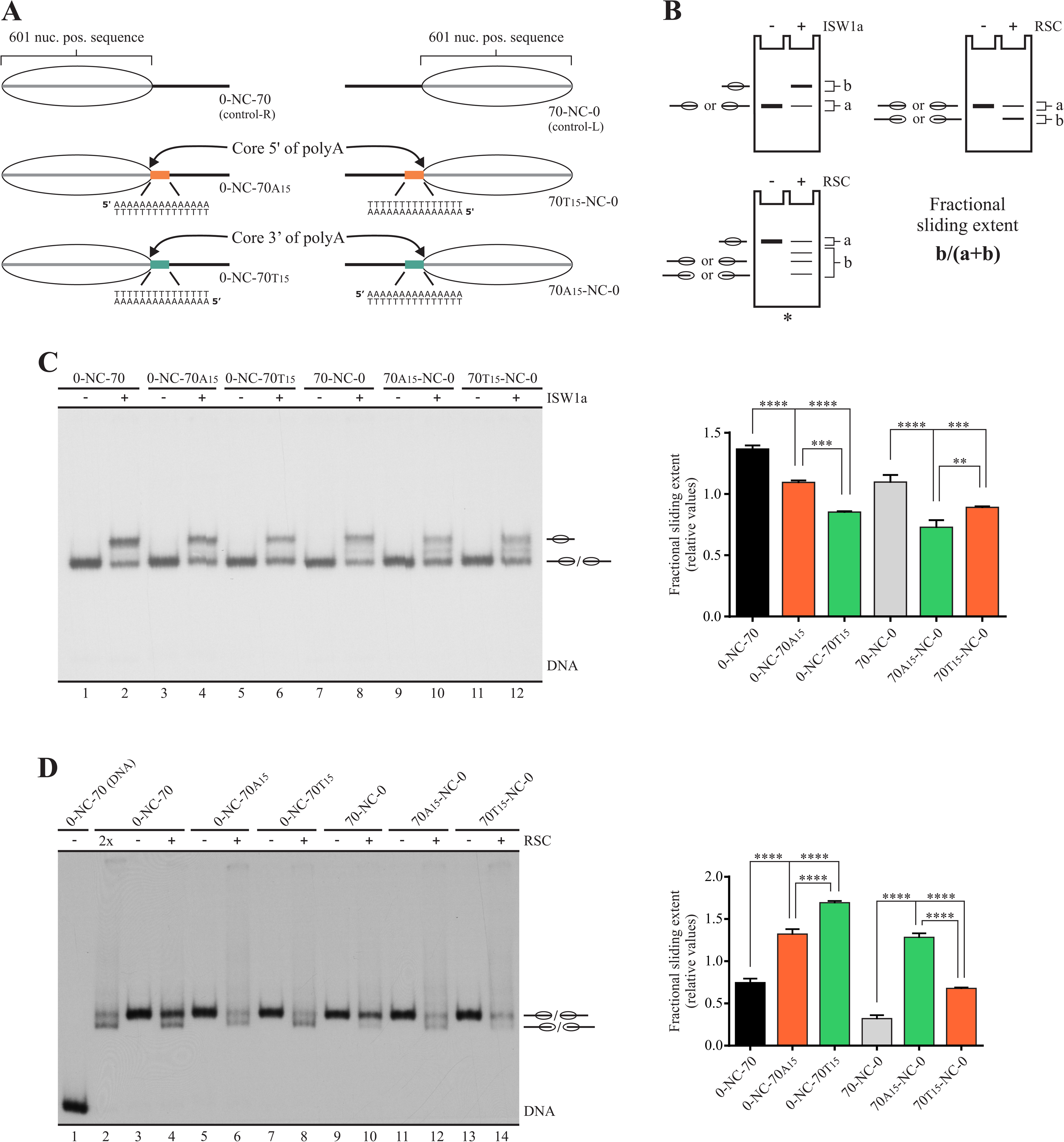
Poly(dA:dT) tracts differentially affect the nucleosome sliding activity of ISW1a and RSC complexes, with the extent of the effects displaying tract orientation-dependency. **(A)** Schematic representation of the nucleosome probes used in the assays. 601 nuc. pos. sequence = nucleosome positioning region of the 601 sequence (gray bar; 147 bp in control probes; 145 bp in tract-containing probes). The oval represents the translational position adopted by the nucleosome core upon reconstitution, which covers the 601 region. The presence and orientation of poly(dA:dT) tracts is represented by colored bars, with the orange bar representing the orientation “core 5’ of polyA” and the green bar representing the orientation “core 3’ of polyA” (see Fig S1 for detailed definitions regarding tract orientations). The term “NC” in probe names stands for “nucleosome core”. Probe names indicate length of linker DNA upstream (left) and downstream (right) the NC and presence of a poly(dA:dT) tract using the letter “A” or “T”, which indicate the sequence of the tract harbored by the upper strand. The number in subscript besides A or T indicates tract length. (B) Schematic representation depicting the remodeling patterns generated by ISW1a and RSC, together with the method used to quantify their activity. The “fractional sliding extent” corresponds to the ratio of intensity given by bands reflecting remodeled (slid) nucleosome over the intensity given by all bands of the nucleosome probe in the lane. The asterisk indicates a type of probe used in assays presented in other figures (nucleosome harboring the core in a central position). (C, D) Nucleosome remodeling assays visualized by electrophoresis in non-denaturing polyacrylamide gels, testing the effect of poly(dA:dT) tracts on ISW1a (C) and RSC (D) complexes. The probe used in each reaction is depicted at the top of each gel picture, where absence or presence of the corresponding remodeling complex is also depicted (1.5 nM ISW1a, 2 nM RSC). Each image is representative of three independent assays, performed under the same conditions. Migrations of alternative forms of the nucleosome probe, which correspond to different translational positions of the histone octamer, are indicated schematically at the right of the picture, where an oval located slightly out of a line indicates migration of a remodeled state generated by RSC that involves mobilization of the histone octamer beyond DNA ends (see text for details). 2x: Remodeling complex concentration two times the concentration used in the other reactions. The graph besides each gel picture depicts quantification of fractional sliding extent from the corresponding assay. Bars in the graphs display the average of three independent assays for each condition analyzed (n = 3). Error bars represent one standard deviation. Asterisks denote statistically significant differences (**p<0.01; ***p<0.001; ****p<0.0001), as deducted from ANOVA with Tukey’s multiple comparisons test.

In order to confirm these effects of poly(dA:dT) tracts, we performed the same analysis on probes harboring linker DNA and the poly(dA:dT) tract on the other side of the 601 sequence (70-NC-0 probes, Fig 1A, right panel). Here, it is worth to remember that the nucleosome has a two-fold symmetry axis (Luger *et al*, 1997; Davey *et al*, 2002). Hence, opposite tract orientations-in terms of presence of a homopolymeric sequence in one strand or the other-correspond to equivalent orientations in terms of nucleosome structure, when one tract is upstream the dyad and the other is downstream this axis. Considering this, an structurally more comprehensive way to describe the two possible orientations of poly(dA:dT) tracts corresponds to “core (or dyad) located on the 5’ side of polyA” and “core (or dyad) located on the 3’ side of polyA” (Fig S1). Thus, we reasoned that, in this new set of probes, the strongest stimulatory effect on RSC sliding activity (and the strongest inhibitory effect on ISW1a activity) should be given by polyA in the upper strand rather than in the lower strand. Of note, the 147 bp 601 sequence is not palindromic; in addition, in Widom’s original 601 plasmid vector as well as in our derivatives, the DNA sequence upstream the 601 positioning region is different to that present downstream this region. To this respect, it has been observed that the 601 sequence bias the directionality of RSC sliding activity (Ghassabi Kondalaji & Bowman, 2022). Thus, the analysis using probes harboring linker DNA upstream the 601 sequence was also designed to ascertain whether poly(dA:dT) tracts exert their influence on RSC and ISW1a activity overcoming these differences between upstream and downstream DNA sequence relative to the 601 axis. The results of the assays performed with this new set of probes confirmed the stimulatory effect of poly(dA:dT) tracts on RSC sliding activity, as well as the inhibitory effect on ISW1a sliding activity. Importantly, as expected from the two-fold symmetry axis of the nucleosome, now the strongest effect (stimulatory for RSC and inhibitory for ISW1a) was given by the presence of polyA in the upper strand (Figs 1C and 1D). Thus, considering both sets of probes, the tract orientation that exerts the strongest stimulatory effect on RSC sliding activity (and the strongest inhibitory effect on ISW1a activity) corresponds to the core located 3’ of polyA. Although differences in the extent of sliding activity were observed when comparing the control probe harboring linker DNA downstream the core to the control harboring linker upstream the core, it did not affect the differential pattern given by tract orientation.

It has been previously shown that poly(dA:dT) tracts located within the nucleosome core stimulate the remodeling activity of RSC (Lorch *et al*, 2014). Considering this earlier finding, we tested the effect of a 15 bp tract whose midpoint is placed on superhelix location (SHL) 5.5 (Fig S2A). By placing a poly(dA:dT) tract on this location we observed no effect on nucleosome stability, relative to the original 601 sequence, in terms of extent of nucleosome reconstitution (see below). In this set of probes, sliding activity of ISW1a remained mostly unaltered (Fig S2B). On the other hand, a stimulatory effect on RSC sliding activity was observed, although this effect was less pronounced than that found on probes harboring the tract on linker DNA and there were no differences between the two alternative tract orientations (Fig S2C).

We next searched for a minimal tract length still having an impact on RSC and/or ISW1a sliding activity, considering that an inverse correlation between the length of poly(dA:dT) tracts and nucleosome occupancy has been reported in previous studies (Field *et al*, 2008). In addition, *in vivo* and *in vitro* studies have found a nucleosome depletion effect of poly(dA:dT) tracts when testing lengths starting from 5 bp (de Boer & Hughes, 2014; Krietenstein *et al*, 2016). First, 15, 10 and 5 bp lengths were tested (Fig 2A). Our results showed that the inhibitory effect on ISW1a sliding activity, as well as the stimulatory effect on RSC activity, were the same when comparing 15 bp to 10 bp length. However, when reducing the length of the tract to 5 bp, the effect was lost for both complexes (Figs 2B and 2C). We then tested the effect of a 7 bp tract, finding an inhibitory effect on ISW1a activity, although weaker than that given by 15 bp length. In the case of RSC, a slight, non-statistically significant, stimulatory effect was observed by 7 bp tract length over control probe (Fig S3). Next, we analyzed the effect of placing progressively longer DNA stretches (0, 19 and 40 bp) between the nucleosome core and a 15 bp poly(dA:dT) tract. To this respect, interactions with linker DNA have been described for both ISW1a and RSC complexes, affecting their interaction with the nucleosome and their remodeling activity (Gangaraju & Bartholomew, 2007; Ye *et al*, 2019; Wagner *et al*, 2020). As observed in Fig 2E, in the probes harboring a separation of 19 and 40 bp between the core and the tract, the inhibitory effect on ISW1a activity was lost. Indeed, even a slight statistically significant stimulatory effect was observed for the probe harboring a 40 bp separation. The analyses performed for RSC showed no variation in the stimulatory effect when comparing 0 and 19 bp separations. In the case of the probe harboring a 40 bp separation between the core and the tract, there was still stimulation of RSC activity, although weaker than that observed for shorter separations (Fig 2F).

**Figure 2.**
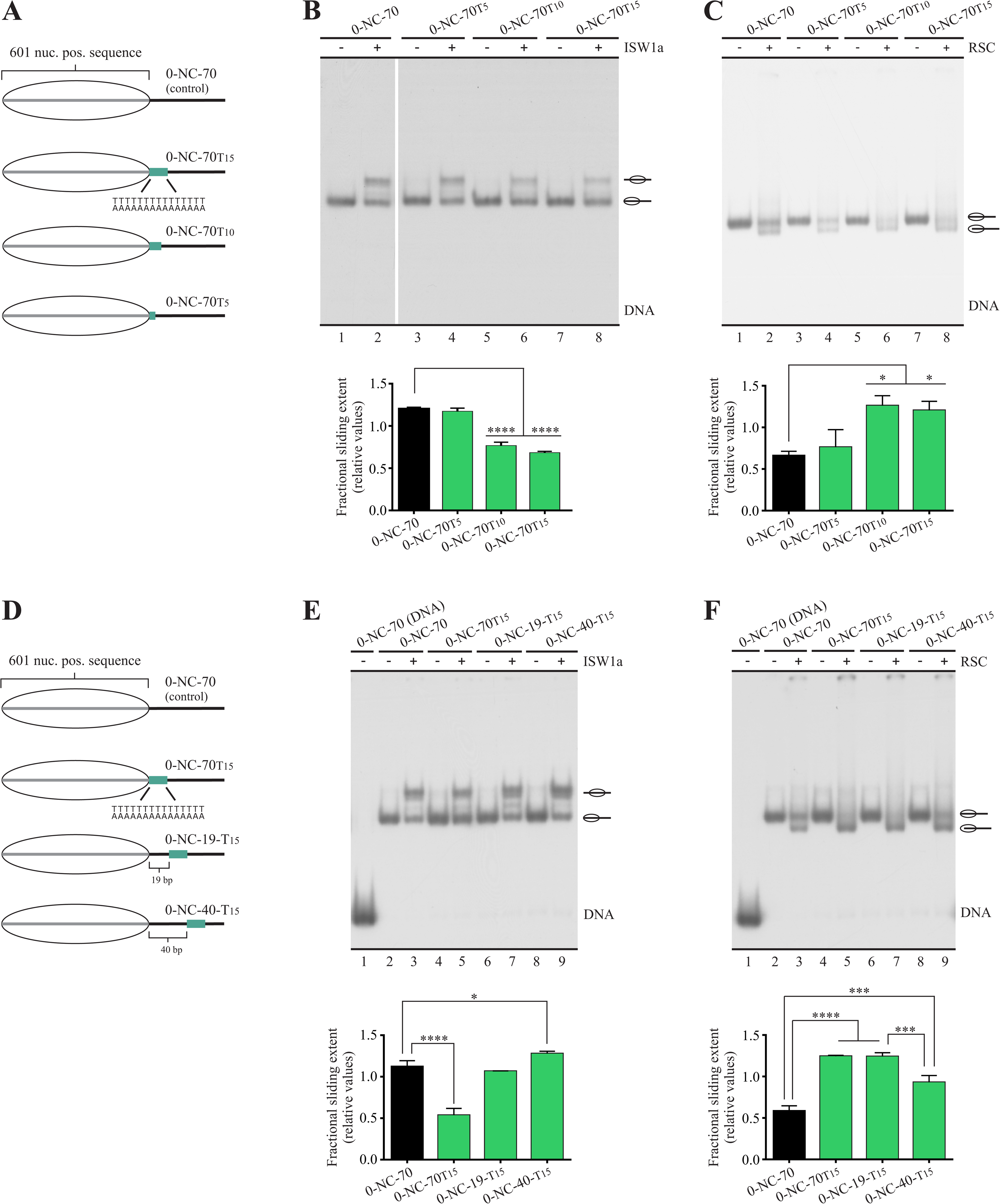
The influence of a poly(dA:dT) tract on nucleosome sliding is affected by its length and particular location in linker DNA. **(A)** Schematic representation of the nucleosome probes used in (B) and (C). **(B, C)** Nucleosome remodeling assays visualized by electrophoresis in non-denaturing polyacrylamide gels, testing the effect of changing tract length on its influence on ISW1a (B) and RSC (C) sliding activity. **(D)** Schematic representation of the nucleosome probes used in (E) and (F). **(E, F)** Nucleosome remodeling assays visualized by electrophoresis in non-denaturing polyacrylamide gels, testing the effect of changing the location of a 15 bp poly(dA:dT) tract on its influence on ISW1a (E) and RSC (F) sliding activity. In figures (B), (C), (E) and (F), the probe used in each reaction is depicted at the top of each gel picture, where absence or presence of the corresponding remodeling complex is also depicted (1.5 nM ISW1a, 2 nM RSC). Each image is representative of three independent assays, performed under the same conditions. Migrations of alternative forms of the nucleosome probe, which correspond to different translational positions of the histone octamer, are indicated schematically at the right of each picture. The graph at the bottom of each gel picture depicts quantification of sliding extent from the corresponding assay (see Fig 1B for information regarding quantification strategy). Bars in the graphs display the average of three independent assays for each condition analyzed (n = 3). Error bars represent one standard deviation. Asterisks denote statistically significant differences (*p<0.05; ***p<0.001; ****p<0.0001), as deducted from ANOVA with Tukey’s multiple comparisons test.

### Poly(dA:dT) tracts exert stronger enhancement of RSC nucleosome disassembling activity when the nucleosome core is located 5’ of polyA

In order to get insight into the mechanisms underlying the differential stimulatory effect given by the two alternative orientations of poly(dA:dT) tracts, we analyzed the influence of these sequences on the strength of RSC-nucleosome interaction. To do this, we performed electrophoretic mobility shift assays (EMSA) using 0-NC-70 probes, previously used to test RSC and ISW1a activity (Fig 1). The effect on the strength of ISW1a-nucleosome interaction was also tested. For these assays, complex concentrations higher than those used in previous remodeling assays were used, in order to obtain observable binding levels. Binding strength was tested in the presence of ATP or in the presence of the non-hydrolyzable analog ATP-γ-S. As expected from previous studies (Wittmeyer *et al*, 2004; Yan *et al*, 2019), the presence of ATP enhanced binding strength of both complexes. This enhancement was observed in the absence and in the presence of poly(dA:dT) tracts (Figs 3A and 3B). Binding of ISW1a to the mononucleosome probe was only detected in the presence of ATP, and no major differences were observed between presence and absence of tracts (Fig 3A). On the other hand, binding of RSC to nucleosome probes harboring a tract was significantly stronger than that observed for the control probe under both conditions, presence of ATP-γ-S (Fig 3B, compare lane 2 to 5 and 8) and presence of ATP (Fig 3B, compare lane 3 to 6 and 9). Interestingly, RSC displayed a higher interaction strength for the probe harboring the core 5’ of polyA than for the probe harboring the core 3’ of polyA (Fig 3B, compare lane 6 to 9). Binding assays performed using the same probes at the form of naked DNA showed stronger binding of RSC to probes harboring a tract, as compared to the absence of tract, with no differences in binding strength between the two tract orientations (Fig S4), indicating that the orientation-dependent differential binding strength arises in the context of the nucleosome structure. At a first glance, the stronger interaction of RSC with the probe harboring the core 5’ of polyA was not consistent with the stimulation of its sliding activity given by the tracts, which was stronger for the probes harboring core 3’ of polyA than for the other orientation (Fig 1). However, electrophoretic analysis in native gels performed in several nucleosome remodeling studies have shown that incubation of mononucleosome probes with RSC or other complexes of the SWI/SNF subfamily results in reduction of the total signal of the mononucleosomal species, accompanied by the appearance of smear or discrete bands reflecting biochemical outcomes such as nucleosome dimers (Lorch *et al*, 1998; Schnitzler *et al*, 1998; Lorch *et al*, 2001; Krajewski & Vassiliev, 2010). It has been proposed that the appearance of these remodeled species corresponds to nucleosomes trapped in structures corresponding to unfinished octamer transfer (Lorch *et al*, 2001; Becker & Horz, 2002). In this context, we reproducibly observed generation of smear and appearance of signal in the loading well in a higher extent for probes harboring the orientation corresponding to core 5’ of polyA. Indeed, when considering the smear signal as part of the remodeled species generated by RSC action, the two alternative tract orientations displayed stimulation of RSC activity at more similar levels (Fig S5). Taken these aspects into account, we reasoned that the stronger binding of RSC to probes harboring core 5’ of polyA might be connected to a stronger octamer transfer activity of this complex, which could result in nucleosome probe disassembling (aka nucleosome eviction). This outcome would be reflected by the appearance of free DNA probe. To test this possibility, we performed remodeling assays using the same high concentrations of RSC used in the binding assays, analyzing the level of nucleosome disassembling obtained for the two alternative tract orientations. Two sets of probes harboring linker DNA downstream the core were tested, one set harbors 70 bp linker DNA (0-NC-70 probes) and the other 30 bp (0-NC-30 probes). For both probe sets, a higher level of nucleosome disassembling was observed for the core located 5’ of polyA, comparing to probes harboring no tract and to probes harboring the other tract orientation (Figs 3C and 3D). In 0-NC-70 probes, the orientation corresponding to core 3’ of polyA resulted in a level of nucleosome disassembling similar to that found in the absence of tract, while in 0-NC-30 probes this orientation resulted in a level of disassembling higher to that observed in the absence of tract. However, this disassembling level was still lower than that found for the orientation corresponding to core 5’ of polyA (Fig 3D, compare lane 7 to 10). Taken together, these results show that a poly(dA:dT) tract exerts a stimulatory effect that is stronger under the orientation corresponding to core located 5’ of polyA than under the opposite orientation, in terms of stimulation of RSC binding to the nucleosome and eviction activity.

**Figure 3.**
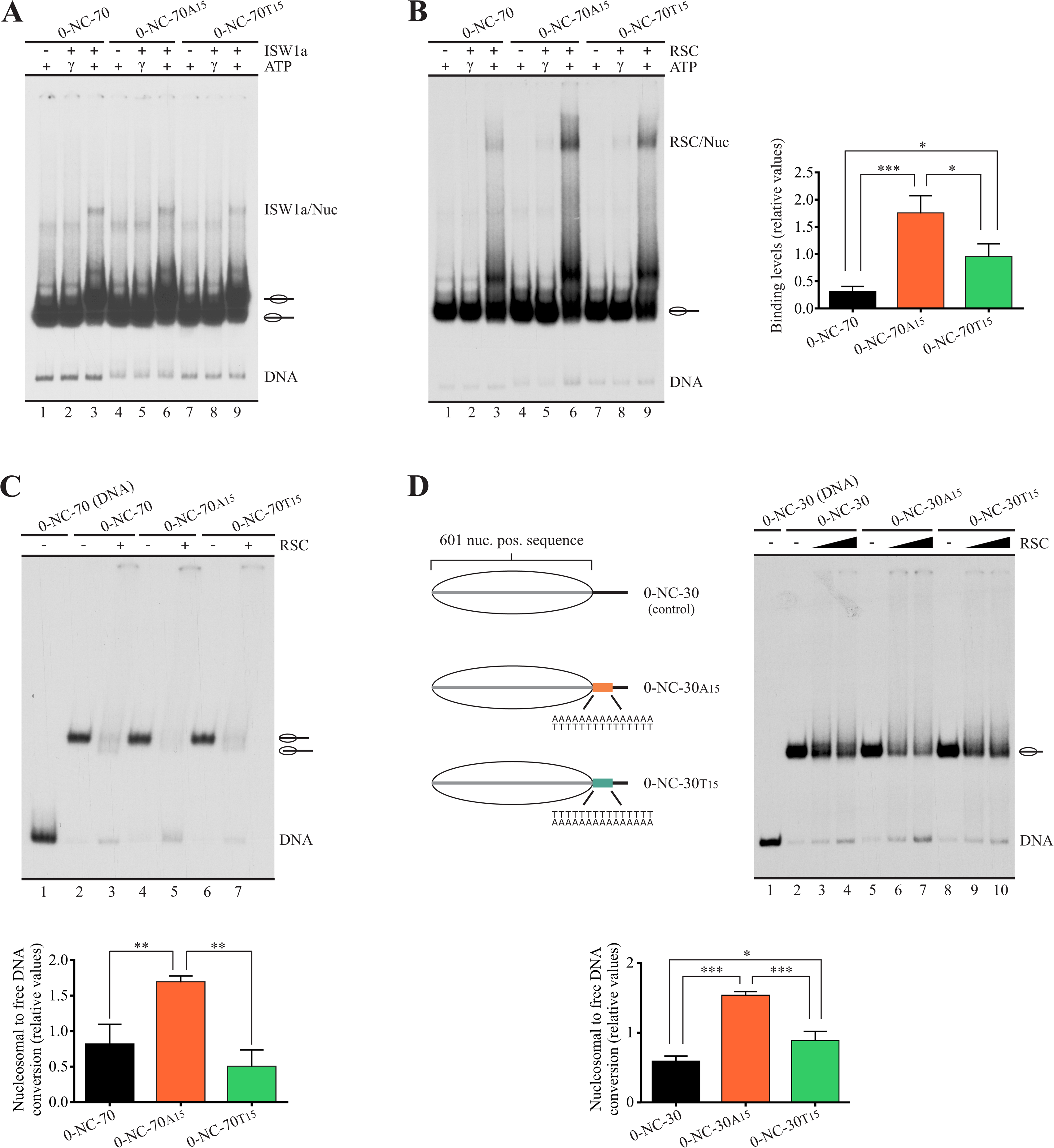
The orientation “core 5’ of polyA” of a poly(dA:dT) tract exerts stronger enhancement of RSC nucleosome disassembling activity than the opposite orientation. **(A-C)** Assays performed using probes harboring a 70 bp linker downstream the nucleosome core, visualized by electrophoresis in non-denaturing polyacrylamide gels. See Fig 1A for a schematic representation of these probes. The assays shown in (A) and (B) correspond to analysis of binding to nucleosomal probes by ISW1a (A) and RSC (B) complexes. The probe used in each reaction is depicted at the top of each gel picture, where absence or presence of ATP, ATP-γ-S and the corresponding remodeling complex is also depicted (9 nM ISW1a, 8 nM RSC). Each gel image is representative of three independent assays, performed under the same conditions. Migrations of the different species is indicated at the right of each picture. The right panel in (B) corresponds to quantification of binding levels of RSC, calculated as a ratio of the bound nucleosomal probe over the total nucleosome probe signal in the lane. Bars in the graph display the average of three independent assays for each condition analyzed (n = 3). Error bars represent one standard deviation. Asterisks denote statistically significant differences (*p<0.05; ***p<0.001), as deducted from ANOVA with Tukey’s multiple comparisons test. The gel image shown in (C) corresponds to a nucleosome remodeling assay performed for RSC (8 nM). The probe used in each reaction is depicted at the top of the gel picture, where absence or presence of the remodeling complex is also depicted. The image is representative of three independent assays, performed under the same conditions. Migrations of the nucleosome probe is indicated schematically at the right of the picture. The graph at the bottom of the gel image depicts quantification of eviction extent from the assay. This was calculated by subtracting the free DNA signal of the probe present in the absence of RSC from the free DNA signal obtained in the presence of this complex; the resulting value was divided by the nucleosomal DNA signal present in the absence of complex, obtaining the eviction extent (fraction of nucleosomal DNA that is converted to free DNA). Bars in the graph display the average of three independent assays for each condition analyzed (n = 3). Error bars represent one standard deviation. Asterisks denote statistically significant differences (**p<0.01), as deducted from ANOVA with Tukey’s multiple comparisons test. **(D)** Nucleosome remodeling assay performed for RSC and probes harboring a short linker DNA downstream the nucleosome core, visualized by electrophoresis in a non-denaturing polyacrylamide gel. *Left panel*: Schematic representation of the nucleosome probes used in the assay. *Right panel*: Nucleosome remodeling assay, performed as described for the assay shown in (C). Low and high RSC concentrations correspond to 2 and 8 nM, respectively. *Bottom panel*: Quantification of eviction extent corresponding to 8 nM RSC, performed as described for the assay shown in (C). Bars in the graph display the average of three independent assays for each condition analyzed (n = 3). Error bars represent one standard deviation. Asterisks denote statistically significant differences (*p<0.05; ***p<0.001), as deducted from ANOVA with Tukey’s multiple comparisons test.

### In a nucleosome harboring linker DNA on both sides of the core but a poly(dA:dT) tract on only one linker, RSC preferentially selects this linker as exit DNA for sliding

As mentioned in the Introduction, detailed structural analyses have recently characterized a module in RSC responsible for interaction with linker DNA (DIM) upon binding of this complex to the nucleosome. Thus, the presence of linker DNA on only one side of the nucleosome core drives directional binding of RSC, making this DNA entry/exit site of the nucleosome the exit DNA in the sliding process catalyzed by the complex. In light of this property of RSC, it has been proposed that this complex interacts with linker DNA that is turned into an NFR at gene promoters, allowing the complex to push the-1 or +1 nucleosomes away from the center of the NFR being generated or expanded (Ye *et al*, 2019; Wagner *et al*, 2020). However, these studies did not analyze what differential properties present on NFR’s DNA favor a preferential binding of the DIM to this linker over the linker DNA located on the other side of the nearest nucleosome core. Properties such as the presence of binding sequences characterized for RSC subunits (Badis *et al*, 2008) and a longer length of the linker/NFR to be extended, relative to upstream and downstream linkers before RSC arrival, would be among differential properties driving directional binding of RSC. In this context, we asked whether the presence of a poly(dA:dT) tract in only one linker of a mononucleosome harboring the same length of linker DNA on both sides, could drive directional sliding activity of RSC. To answer this question, we performed sliding assays using probes harboring 35 bp of linker DNA on both sides of the nucleosome core (35-NC-35) and a poly(dA:dT) tract located right upstream or downstream the core, assessing directionality of sliding activity by digestion with restriction enzymes after the remodeling step and after removal of the complex (Fig 4A). The two alternative tract orientations were tested for probes harboring tract upstream the core as well as for probes harboring tract downstream the core (Fig 4B). As in our previous analyses performed on probes harboring linker DNA on only one side of the core, the presence of tracts stimulated the sliding activity of RSC, with probes harboring the core 3’ of polyA giving the strongest stimulation, in terms of slid over total mononucleosome signal (Fig S6).

**Figure 4.**
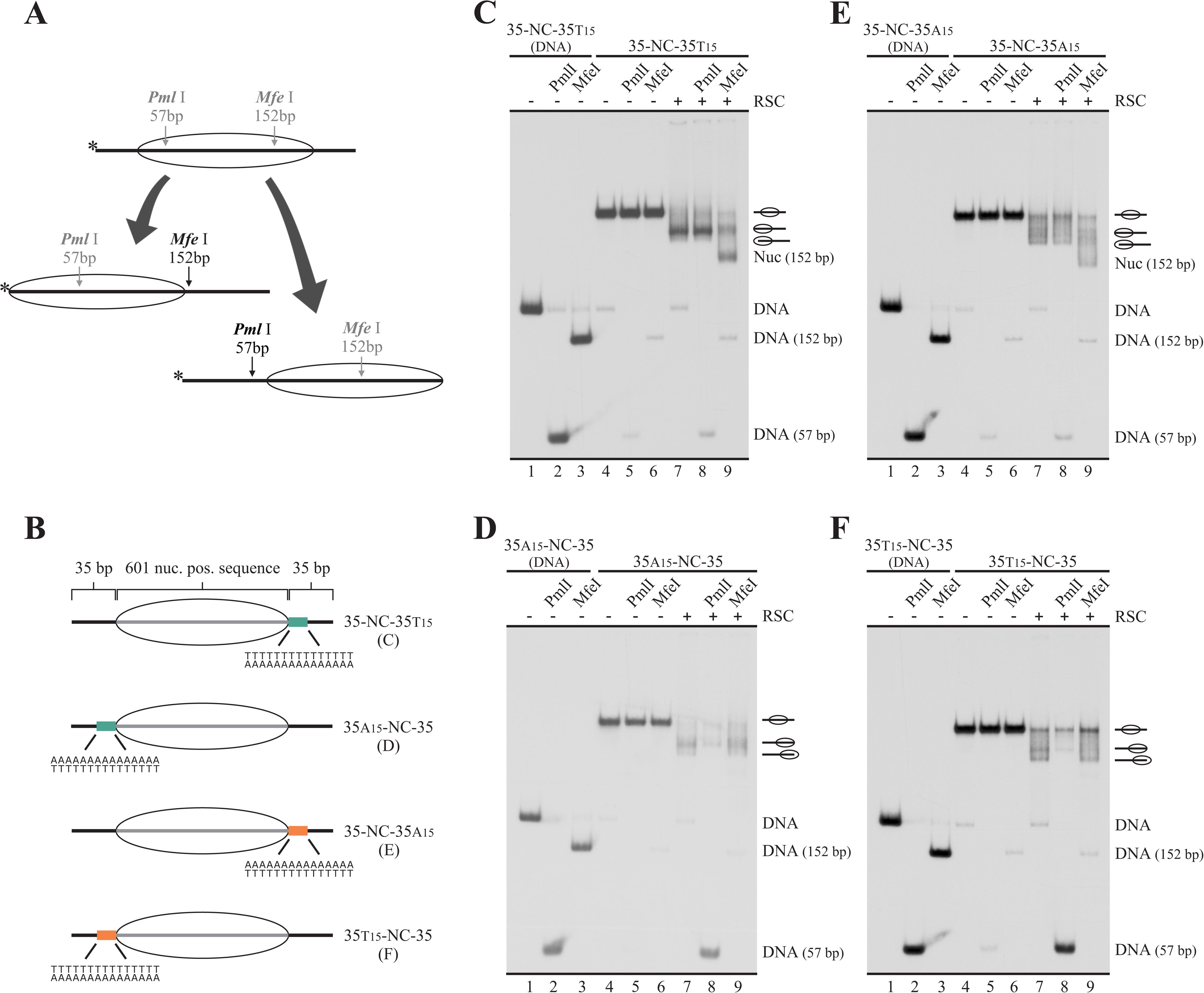
On nucleosomes harboring same-length linkers on both sides of the core, RSC selects the tract-containing linker as the exit DNA. **(A)** Schematic representation depicting changes in accessibility to restriction enzymes depending on upstream or downstream mobilization of the histone octamer. Names of restriction enzymes presented in gray represent absence of accessibility, while names presented in black represent access of the corresponding restriction site. The numbers below the names of restriction enzymes indicate the size of labeled DNA segments resulting from digestion by the corresponding enzyme, relative to the 5’ end of the upper strand (labeled DNA end, represented by an asterisk). Note that cutting of the nucleosomal probe by *Pml*I results in a labeled fragment corresponding to naked DNA (57 bp), while cutting of the nucleosomal probe by *Mfe*I results in a 152 bp labeled fragment corresponding to a nucleosome. **(B)** Schematic representation of the nucleosome probes used in the assays. **(C-F)** Nucleosome remodeling assays testing directionality of RSC sliding activity by changes in accessibility to restriction enzymes, visualized by electrophoresis in non-denaturing polyacrylamide gels. The probe used in each assay is depicted at the top of each gel picture, where the presence of RSC (2 nM) and/or a defined restriction enzyme is also depicted. Each image is representative of three independent assays, performed under the same conditions. Migration of different species is depicted at the right of each gel picture. In all assays, lanes 1 to 3 correspond to probe at the form of naked DNA (mock reconstitution).

We first tested a probe harboring the tract downstream the core, with polyA in the lower strand (35-NC-35_T15_). As observed in Fig 4C, on this probe RSC mobilized the histone octamer mainly upstream, reflected by minimal digestion of the slid nucleosome by *Pml*I (cut probe signal mainly coming from the small fraction of naked DNA present in the probe) and strong digestion by *Mfe*I (Fig 4C, compare lane 8 to 9). We then tested a probe harboring the tract upstream the core, with polyA in the upper strand (35_A15_-NC-35). Using this probe, RSC mobilized the histone octamer mainly downstream, reflected now by strong digestion of the slid nucleosome by *Pml*I and minimal digestion by *Mfe*I (Fig 4D, compare lane 8 to 9). These results demonstrate that the presence of the poly(dA:dT) tract determines the selection of the exit DNA for the sliding activity catalyzed by RSC, ensuring mobilization of the histone octamer away from the tract. Importantly, the same results were obtained when testing the opposite tract orientation (core 5’ of polyA, Figs 4E and 4F), indicating that sliding of the nucleosome core away from the tract by RSC is independent of tract orientation. This effect of poly(dA:dT) tracts on RSC sliding activity was also confirmed using mononucleosome probes harboring linker DNA on only one side of the core (Fig S7).

Regarding the effects of poly(dA:dT) tracts on directionality of sliding activity, we asked whether these tracts hinder ISW1a activity when they are not located in the way of the histone octamer movement during the sliding reaction catalyzed by this complex. In this regard, it has been demonstrated that ISW1a is highly sensitive to the presence and length of linker DNA and that this complex mobilizes the histone octamer towards the linker DNA to which it is bound (Gangaraju & Bartholomew, 2007). In the case of *in vitro* assays performed using mononucleosome probes harboring linker DNA on only one side of the nucleosome core, the complex mobilizes the histone octamer towards a central position in the DNA segment. Thus, in this type of probe, ISW1a attempts to mobilize the histone octamer towards the poly(dA:dT) tract if this sequence is present in the linker DNA, as was the case of our initial analyses performed using nucleosome probes harboring a 70 bp linker DNA on only one side of the core (Figs 1, 2 and S3). In order to assess whether poly(dA:dT) tracts still inhibit ISW1a activity when located on linker DNA that becomes exit DNA upon ISW1a engagement, we tested probes harboring a long linker upstream the core and a tract-containing short linker downstream the core (70-NC-30 probes, Fig 5A). Under this design, poly(dA:dT) tracts did not hinder ISW1a sliding activity; In fact, there was a slight stimulation of its activity (Figs 5B and 5C). The same result was obtained using a probe harboring the shorter linker and a tract upstream the nucleosome core (Fig S8).

**Figure 5.**
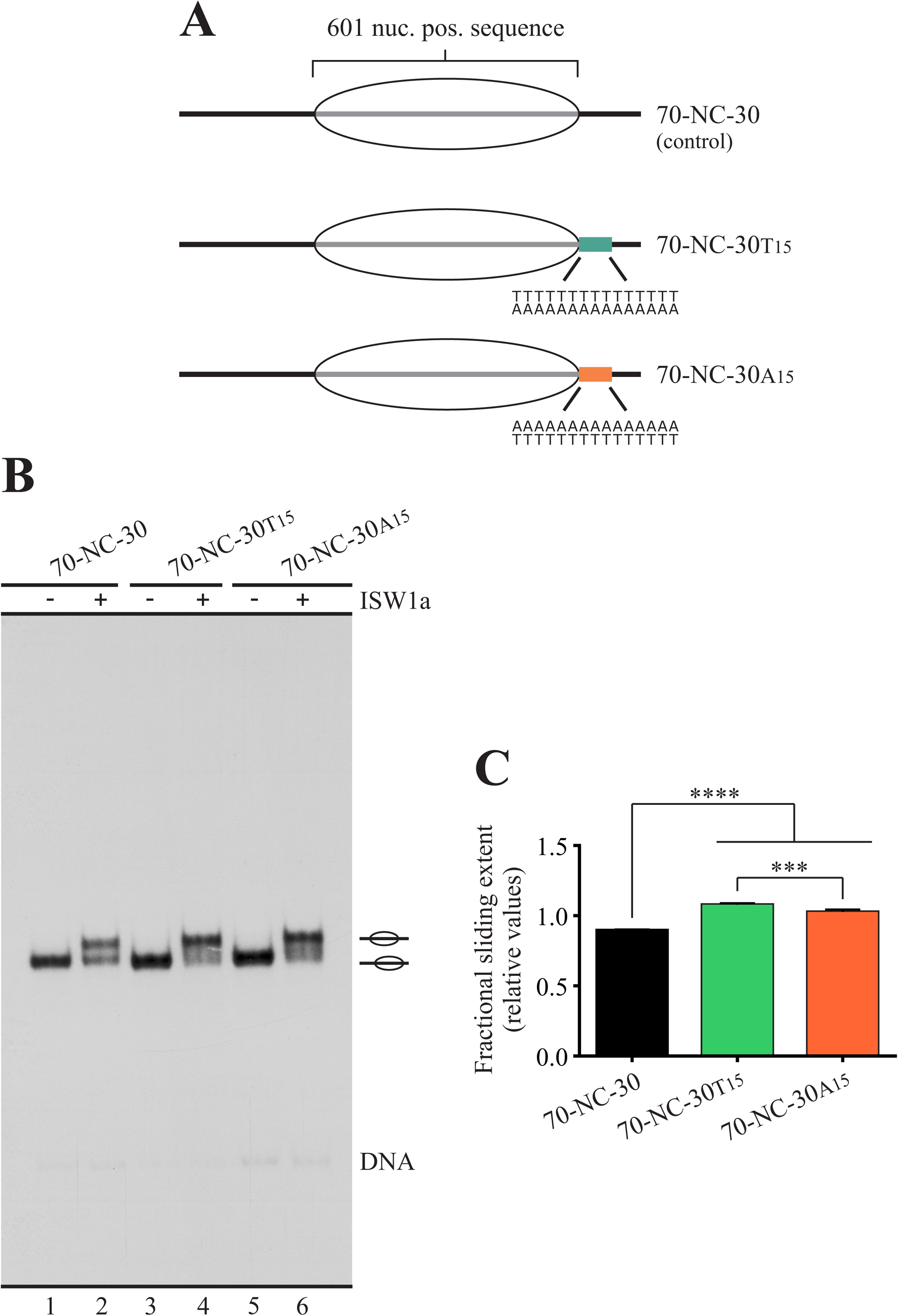
Poly(dA:dT) tracts do not hinder ISW1a activity when located in linker that becomes exit DNA during the sliding process. **(A)** Schematic representation of the nucleosome probes used in the assay. **(B)** Nucleosome remodeling assay visualized by electrophoresis in a non-denaturing polyacrylamide gel, testing the effect of a poly(dA:dT) tract, located in the shorter linker, on ISW1a sliding activity. The probe used in each reaction is depicted at the top of the gel picture, where absence or presence of the remodeling complex (1.5 nM) is also depicted. The image is representative of three independent assays, performed under the same conditions. Migrations of alternative forms of the nucleosome probe, which correspond to different translational positions of the histone octamer, are indicated schematically at the right of the picture. **(C)** Quantification of sliding extent (see Fig 1B for information regarding quantification strategy). Bars in the graph display the average of three independent assays for each condition analyzed (n = 3). Error bars represent one standard deviation. Asterisks denote statistically significant differences (***p<0.001; ****p<0.0001), as deducted from ANOVA with Tukey’s multiple comparisons test.

### A design that forces RSC to mobilize a tract across the nucleosome core results in blocking of its sliding activity by one tract orientation

Our finding that RSC sliding activity always proceeds by mobilizing the nucleosome core away from the poly(dA:dT) tract, independently of tract orientation, led us to ask what would be the effect of a tract on the sliding activity of RSC if the complex is forced to mobilize the tract across the nucleosome core. To test this scenario, we generated mononucleosome probes harboring linker DNA upstream and downstream the core, with both linkers being of the same length and both harboring a 15 bp poly(dA:dT) tract (35-NC-35 probes, Figs 6A and S9A). We first analyzed a probe harboring the tract orientation that results in the strongest stimulatory effect on RSC sliding activity, corresponding to the core located 3’ of polyA (35_A15_-NC-35_T15_, Fig S9A). As expected, the presence of tracts on both sides of the core resulted in a stronger stimulatory effect of RSC sliding activity, as compared to the presence of only one tract (Fig S9). Intriguingly, RSC action on this probe did not result in generation of the fastest migrating nucleosomal band. However, its action did result in generation of this remodeled species in centrally positioned nucleosomes harboring either one tract (Fig S9) or two tracts in the opposite orientation (core 5’ of polyA, 35_T15_-NC-35_A15_, Fig 6A), and even in the control probe under longer incubations (Fig S9D). As mentioned above, it has been previously determined that the remodeled state present in this fastest migrating band corresponds to a nucleosome harboring a histone octamer slid up to 51 bp beyond a DNA end, resulting from the action of RSC; this outcome extends to other complexes of the SWI/SNF subfamily (Flaus & Owen-Hughes, 2003; Kassabov *et al*, 2003). Other studies have also found a fastest migrating band reflecting loss of one H2A-H2B heterodimer (Bruno *et al*, 2003; Lorch *et al*, 2006), an effect that would also involve mobilization of the histone octamer beyond a DNA end (Bruno *et al*, 2003). Thus, in light of the result obtained in the sliding assays testing the probes harboring poly(dA:dT) in both linkers, we reasoned that there might be a region inside the nucleosome core where the presence of a tract in one specific orientation (dyad 3’ of polyA) could make sliding activity of this complex more difficult. Thereafter, in a probe harboring longer tract-containing linkers, with both tracts in the orientation core 3’ of polyA, the sliding reaction would not only result in the absence of the fastest migrating band, but even in the absence of bands reflecting relocation of the histone octamer from the starting central position to positions reaching DNA ends. To test this prediction, we performed sliding assays using 70-NC-70 probes (Fig 6B). Under this configuration, where each tract is 2 bp inside the core after nucleosome reconstitution and there are 70 bp for histone octamer mobilization in *cis* until reaching a DNA end, octamer mobilization reaching these ends forces that one of the tracts travels all the way up to the nucleosome dyad. As predicted, RSC action on the probe harboring the configuration consisting of core located 3’ of each polyA sequence resulted in mobilization of the histone octamer to translational positions that do not reach DNA ends (Fig 6B, lane 4). On the other hand, RSC action on the probe harboring the opposite orientation in both tracts did result in relocation of the core to translational positions reaching DNA ends (Fig 6B, lane 6). For this probe, at least two faster-migrating bands are obtained, relative to the main band obtained for the probe harboring the orientation core 3’ of polyA (Fig 6B, compare lane 4 to 6).

**Figure 6.**
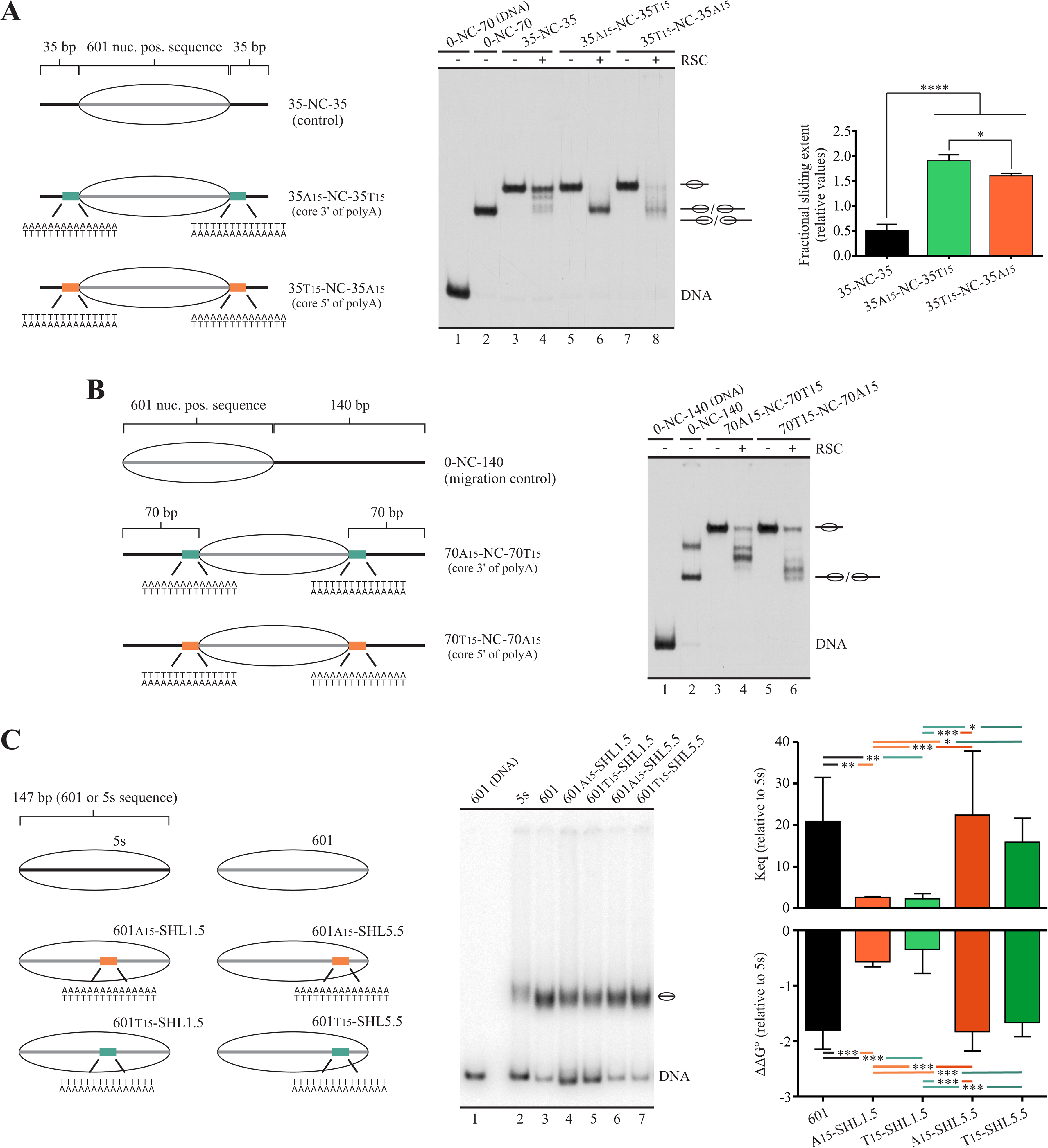
A design that forces RSC to mobilize a tract across the nucleosome core results in blocking of its sliding activity by the orientation “core 3’ of polyA”. **(A)** Remodeling assay using probes harboring short same-length linkers on both sides of the nucleosome core, each containing a poly(dA:dT) tract. *Left panel*: Schematic representation of the nucleosome probes used in the assay. *Central panel*: Nucleosome remodeling assay visualized by electrophoresis in a non-denaturing polyacrylamide gel. The probe used in each reaction is depicted at the top of the gel picture, where absence or presence of the remodeling complex (2 nM) is also depicted. The image is representative of three independent assays, performed under the same conditions. Migrations of alternative forms of the nucleosome probe, which correspond to different translational positions of the histone octamer, are indicated schematically at the right of the picture. *Right panel*: Quantification of sliding extent (see Fig 1B for information regarding quantification strategy). Bars in the graph display the average of three independent assays for each condition analyzed (n = 3). Error bars represent one standard deviation. Asterisks denote statistically significant differences (*p<0.05; ****p<0.0001), as deducted from ANOVA with Tukey’s multiple comparisons test. **(B)** Remodeling assay using probes harboring long same-length linkers on both sides of the nucleosome core, each containing a poly(dA:dT) tract. *Left panel*: Schematic representation of the nucleosome probes used in the assay. *Right panel*: Nucleosome remodeling assay visualized by electrophoresis in a non-denaturing polyacrylamide gel. See legend of central panel in (A) for a description of the remodeling assay. **(C)** Competitive nucleosome reconstitution analysis. *Left panel*: Schematic representation of the nucleosome probes used in the assay. 5s: 147 bp probe corresponding to the dominant positioning region of the *Lytechinus variegatus* 5s rDNA. SHL: superhelix location (1.5 or 5.5). *Central panel*: Competitive nucleosome reconstitution assay visualized by electrophoresis in a non-denaturing polyacrylamide gel. The probe used in each reconstitution reaction is depicted at the top of the gel picture. The image is representative of three independent assays, performed under the same conditions, two in triplicate and one in duplicate (n = 8). Migrations of the probes at the form of naked DNA and reconstituted mononucleosome are indicated at the right of the picture. *Right panel*: Graphs depicting values of equilibrium constant (*K*eq) and ΔΔ*G*° of nucleosome formation, relative to 5s, calculated from the competitive nucleosome reconstitution assay. Error bars represent one standard deviation. Asterisks denote statistically significant differences (*p<0.05; **p<0.01; ***p<0.001), as deducted from ANOVA with Tukey’s multiple comparisons test.

Considering that poly(dA:dT) tracts have been described as stiff DNA sequences that negatively affect nucleosome stability (Anderson & Widom, 2001; Segal & Widom, 2009), we asked whether this property of the tracts would be the factor underlying this particular effect on RSC sliding activity. If so, one tract orientation should display a deeper negative impact on nucleosome stability. To test this possible mechanism, we performed competitive nucleosome reconstitution analyses (Thastrom *et al*, 2004) on 147 bp DNA segments (Fig 6C) harboring a 15 bp tract in one of the two alternative orientations and one of two alternative locations: Midpoint of the tract 15 bp downstream the 601’s dyad (A15-SHL1.5 and T15-SHL1.5) and midpoint 54 bp downstream the 601’s dyad (A15-SHL5.5 and T15-SHL5.5). Regarding this, it has been determined that the regions of higher distortion of DNA in the nucleosome structure might vary with changes in DNA sequence (Cui & Zhurkin, 2010; Vasudevan *et al*, 2010; Frouws *et al*, 2016). In the case of the 601 sequence, the regions of higher distortion of DNA correspond to SHL1.5 and SHL4.5 (Vasudevan *et al*, 2010). Therefore, in one set of probes the tract covers a region that, upon reconstitution, corresponds to one of the points of higher distortion for the 601 sequence, while in the other set the tract locates near but do not overlap with the other region of higher distortion. Our results showed that locating a tract in SHL5.5 has no effect on nucleosome stability, but placement on SHL1.5 resulted in a marked reduction of nucleosome stability, with an 8 to 10 fold reduction in the relative equilibrium constant of nucleosome formation (Fig 6C). Importantly, no differences in reduction of nucleosome stability were found when comparing the two alternative tract orientations in any of the locations tested (Fig 6C), discarding the effects of poly(dA:dT) tracts on nucleosome stability as the mechanism underlying this differential effect exerted by the two alternative tract orientations on RSC sliding activity.

### Poly(dA:dT) tracts globally influence the directionality of ISW1a sliding activity

The results of our analyses testing the effect of poly(dA:dT) on RSC activity are in agreement with the general trend observed both *in vitro* and *in vivo* for single-tract containing NFRs, which shows a longer stretch of the NFR located 5’ of polyA (de Boer & Hughes, 2014), and with analyses performed by Krietenstein and co-workers (Krietenstein *et al*, 2016), which show that RSC displaces nucleosomes asymmetrically around these tracts (see Discussion for further details). In addition, this study shows that ISW1a displaces nucleosomes symmetrically around poly(dA:dT) tracts, with displacement of surrounding nucleosomes away from the tract. However, an analysis of directionality of ISW1a sliding activity in terms of presence versus absence of poly(dA:dT) tracts was not performed. On the other hand, it has been previously demonstrated that the *in vivo* action of ISW1a results in nucleosomes populating gene promoter NFRs (Parnell *et al*, 2015). In this context, the results of our assays testing ISW1a activity predict that ISW1a would act reducing the length of non-tract-containing NFRs and expanding the length of tract-containing NFRs. To test this prediction, we analyzed publicly available data generated by Krietenstein and co-workers, who performed genome-wide reconstitution of the *S. cerevisiae* genome into nucleosomes and determined nucleosome repositioning mediated by several factors and complexes, including ISW1a (Krietenstein *et al*, 2016). From these data, we determined the frequency of polyA and polyT tracts in the sense strand of gene promoters that displayed mobilization of the-1 nucleosome by ISW1a in at least 20 bp. Three clusters were generated: *All genes*, corresponding to the full gene promoter coverage of the study (4,064 genes); *upstream mobilization*, corresponding to gene promoters displaying ISW1a-mediated upstream mobilization of the-1 nucleosome (1,793 genes); and *downstream mobilization*, corresponding to gene promoters displaying ISW1a-mediated downstream mobilization of the-1 nucleosome (854 genes). As the analysis searched for presence of poly(dA:dT) tracts that might influence ISW1a activity, the frequency of poly(dA:dT) tracts was determined relative to the position of the-1 nucleosome found in the absence of ISW1a (termed SGD condition). The same analysis was performed for the +1 nucleosome; in this case, 890 genes displayed ISW1a-mediated upstream mobilization and 1,543 displayed downstream mobilization of this nucleosome. As expected from previous studies, there is a high frequency of poly(dA:dT) tracts right downstream the-1 nucleosome and right upstream the +1 nucleosome under the SGD condition, together with a low frequency near-1 and +1 dyads, which reflects the translational location passively adopted by histone octamers upon reconstitution (Figs 7A and 7B, respectively). Interestingly, in line with our prediction, for those gene promoters that displayed upstream mobilization of the-1 nucleosome by ISW1a, the frequency under SGD condition is higher right downstream the-1 nucleosome core (Fig 7A). Conversely, for the cluster that displayed downstream mobilization of the-1 nucleosome the frequency of poly(dA:dT) tracts under SGD condition is lower right downstream the-1 nucleosome core, relative to the other clusters (Fig 7A). The equivalent result was obtained for the +1 nucleosome. Hence, a low frequency of poly(dA:dT) tracts-relative to the other clusters-was found in the cluster of genes that displayed upstream mobilization of the nucleosome by ISW1a, in the region right upstream the +1 nucleosome core (Fig 7B). Interestingly, a higher frequency of poly(dA:dT) tracts downstream this nucleosome can be observed for this cluster, relative to the frequency observed for the other clusters in that region (Fig 7B). Taking together, our bioinformatics analyses show that in those genes displaying ISW1a-mediated nucleosome mobilization towards the NFR, for both-1 and +1 nucleosomes, there is a lower frequency of poly(dA:dT) tracts in this region.

**Figure 7.**
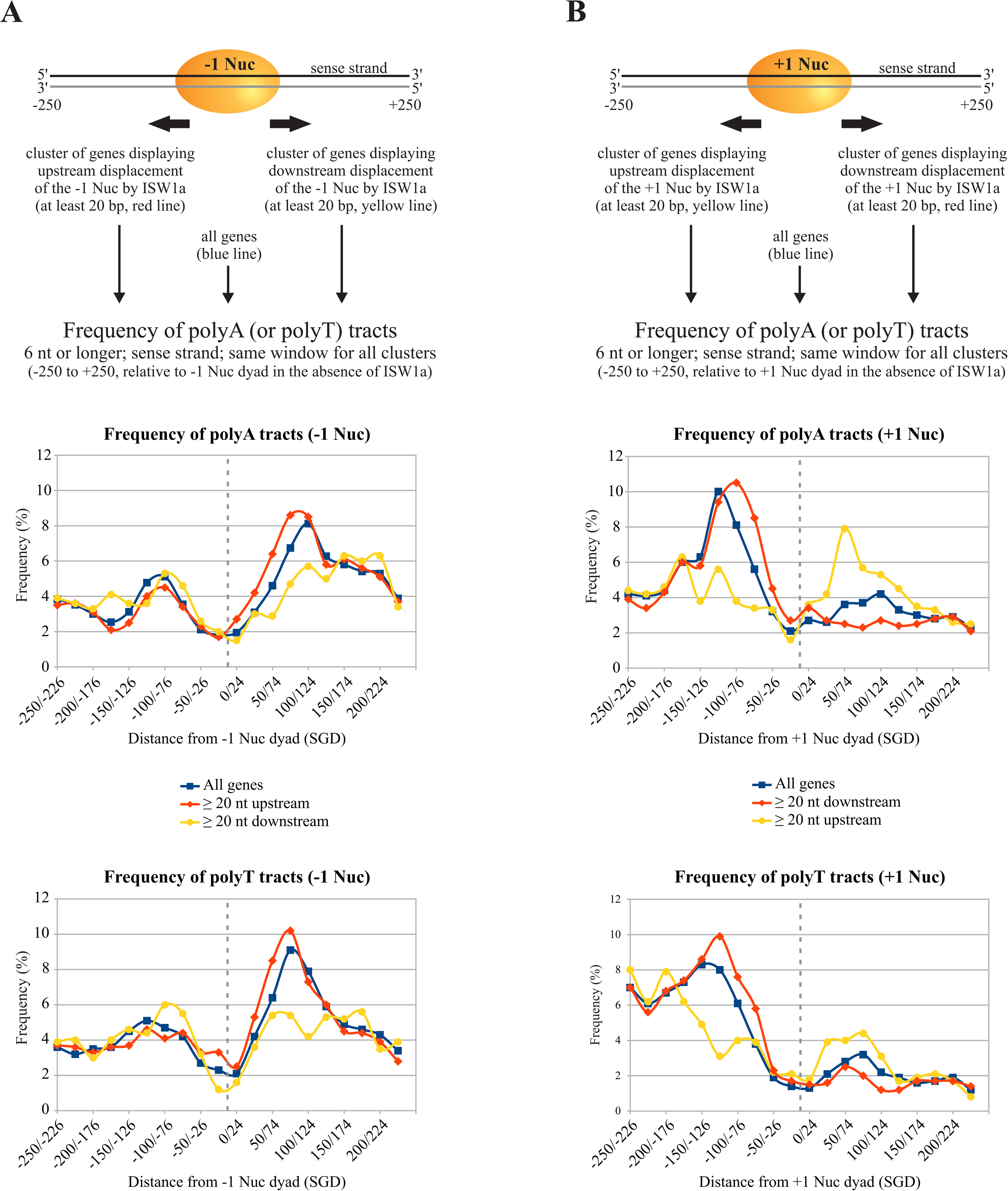
Genes undergoing mobilization of-1 and +1 nucleosomes towards their promoter NFR by ISW1a action, display a low frequency of poly(dA:dT) tracts right downstream and upstream the original position of-1 and +1 nucleosomes, respectively. Bioinformatics analysis of the effects of ISW1a on *in vitro* reconstituted promoter nucleosomes of the *S. cerevisiae* genome. The analysis was performed from data generated by Krietenstein and co-workers (Krietenstein *et al*, 2016). Nucleosome positioning data was taken from two conditions: absence of any CRC or factor (SGD, starting position of-1 and +1 nucleosomes) and treatment with the ISW1a complex. For both conditions, the positions of-1 and +1 nucleosomes in every gene (4064 genes) was determined. Afterwards, the difference in positioning of these nucleosomes (SGD vs ISW1a treatment) was determined for each gene. Subsequently, four clusters were generated: *i*) Genes displaying more than 20 bp upstream displacement of the-1 nucleosome; *ii*) Genes displaying more than 20 bp downstream displacement of the-1 nucleosome; *iii*) Genes displaying more than 20 bp upstream displacement of the +1 nucleosome; and *iv*) Genes displaying more than 20 bp downstream displacement of the +1 nucleosome. Then, the frequency of poly(dA:dT) tracts-binned in 25 bp intervals-was determined for these clusters, as well as for all genes, in a window of 500 bp around the-1 (or +1) nucleosome dyad peak found in the absence of ISW1a (SGD condition). This frequency was calculated as the number of tract-containing genes over the total number of genes in the corresponding cluster (times 100 percent), for each interval. In the graphs, the yellow labels represent the clusters that display nucleosome displacement towards the NFR upon ISW1a action (downstream displacement for-1 Nuc and upstream displacement for +1 Nuc), while red labels represent the clusters that display nucleosome displacement away from the NFR upon ISW1a action (upstream displacement for-1 Nuc and downstream displacement for +1 Nuc). The analysis is schematically described at the top of each figure. **(A)** Analysis performed for-1 nucleosome. **(B)** Analysis performed for +1 nucleosome.

## Discussion

In this work we performed a systematic analysis addressing the effect of poly(dA:dT) tracts on the nucleosome remodeling activity of ISW1a and RSC complexes, finding new properties of these DNA sequences in terms of their influence on ATP-dependent nucleosome remodeling. These properties are: (*i*) tracts enhance RSC binding to nucleosomes and dictate what linker will be the exit DNA in the sliding process, resulting in nucleosome core mobilization away from the tracts; (*ii*) although by different means, these sequences also impose sliding of the nucleosome core away from them to the ISW1a complex; (*iii*) tracts enhance RSC eviction activity when being in one of the two possible orientations relative to a neighboring nucleosome core; and (*iv*) in a scenario where RSC is forced to mobilize a poly(dA:dT) tract inside the nucleosome core, one tract orientation blocks the sliding activity of this complex.

### Poly(dA:dT) tracts exert orientation-dependent and-independent effects on RSC activity

Our findings show that, when a poly(dA:dT) tract is present in linker DNA, one of its orientations (core 5’ of polyA) exert a stronger stimulation on RSC binding to the nucleosome and, consistently, a stronger stimulation of its nucleosome disassembling activity. On the other hand, the alternative orientation (core 3’ of polyA) exerts a strong stimulation of RSC sliding activity. These effects correspond to differential, orientation-dependent, effects exerted by poly(dA:dT) tracts on RSC activity. The non-differential effect is entailed in the stimulation of RSC sliding activity by the tracts. Here we found that RSC slides the histone octamer away from the tracts, independently of their orientation. This result also implies that RSC selects the tract-containing linker as the exit DNA during the remodeling process. In this context, as mentioned in the Introduction, recent structural analyses have characterized a linker DNA-interaction module (DIM) in the complex, with this linker DNA being the exit DNA during nucleosome remodeling (Ye *et al*, 2019; Wagner *et al*, 2020). Considering this property of RSC, our findings suggest that both tract orientations act in the same way on RSC, consisting in a stimulatory effect on DIM binding to linker DNA. Then, the stronger binding stimulation given by one tract orientation (core 5’ of polyA) would make a nucleosome more prone to be evicted. To this respect, single-molecule experiments have observed generation of intranucleosomal loops of DNA during the remodeling process catalyzed by SWI/SNF and RSC (Zhang *et al*, 2006). We speculate that these loops might be generated by accumulation of exit DNA between this border of the nucleosome core and the point where the DIM binds to exit linker DNA (Ye *et al*, 2019; Wagner *et al*, 2020). Then, the presence of a poly(dA:dT) tract on linker DNA in the orientation “core 5’ of polyA” would facilitate the generation of this loop and, in turn, histone octamer eviction as remodeling outcome. Alternatively, or being part of the process, this enhanced binding to linker DNA might result in improving the coupling between DNA translocation and ATP hydrolysis (Clapier *et al*, 2016).

Another differential effect exerted by poly(dA:dT) tracts on RSC activity comes from a design that forces this complex to mobilize a tract across the nucleosome core. Under this special design, that involves the presence of tracts on both borders of the nucleosome core, sliding activity of RSC is initially stimulated and then blocked by the orientation “core 3’ of polyA”. To this respect, our competitive nucleosome reconstitution analyses showed that the two alternative orientations of poly(dA:dT) tracts affect nucleosome stability in the same extent, discarding this property of tracts as the mechanism underlying this effect. In this context, it has been described that the tracking lobes of the catalytic subunit of most remodeling complexes, including RSC, engage at the SHL2 position and perform tracking in one strand, in the 3’ to 5’ direction (Saha *et al*, 2005; Clapier, 2021). This property of RSC and the tract orientation that blocks its sliding activity suggest that for the catalytic subunit of the complex it is more difficult to track on polyA than on polyT and other sequences. Nevertheless, it is worth to recall that this effect results from a special design. In the presence of a single tract in one of the linkers of a nucleosome probe, RSC selects this linker as the exit DNA to exert its remodeling activity, implying that there is no entrance of the tract into the nucleosome core during the remodeling process. Thus, RSC does not require its motor to perform tracking on poly(dA:dT) sequences for stimulation of its sliding and eviction activities.

### Poly(dA:dT) tracts hinder ISW1a activity when located in the way of histone octamer mobilization

As mentioned in the Results section, ISW1a is highly sensitive to the presence and length of linker DNA. Upon binding to the nucleosome, ISW1a preferentially binds to the longer linker and makes this linker (proximal linker) the entry DNA in the sliding process (Gangaraju & Bartholomew, 2007). Our analyses show that, when located on this linker, poly(dA:dT) tracts hinder the sliding activity of this complex. Conversely, when placed on the distal linker, there was an even slight enhancement of its activity. The differential effect given by the two alternative tract orientations was essentially observed in the case of hindering of its sliding activity, obtained when the tracts were located in the proximal linker, where the stronger hindering was given by the orientation consisting in “core 3’ of polyA”. According to our results, this inhibition of ISW1a sliding activity would not be owing to the intrinsic antinucleosomal properties of poly(dA:dT) tracts. Our competitive nucleosome reconstitution analyses showed that poly(dA:dT) tracts do not reduce nucleosome stability when placed near the border of the core and display no orientation-dependent effects on nucleosome stability. In addition, we did not detect an effect of tracts on ISW1a binding to the nucleosome. These results suggest that poly(dA:dT) tracts, when located in the proximal linker DNA, might hinder the productive engagement of ISW1a onto the nucleosome, with the orientation “core 3’ of polyA” being more efficient in generating this effect.

### Effects of poly(dA:dT) tracts on ISW1a and RSC at NFRs

Our findings are in accordance with the known role played by poly(dA:dT) tracts at gene promoter NFRs, consisting in establishment and maintenance of these regions (Struhl & Segal, 2013; Barnes & Korber, 2021), and with the opposite roles played by RSC and ISW1a at these genomic regions, where these complexes aim to widen and shrink these regions, respectively (Parnell *et al*, 2015). In addition, our results support a model for the asymmetric chromatin opening that take place around poly(dA:dT) tracts (Fig 8).

**Figure 8.**
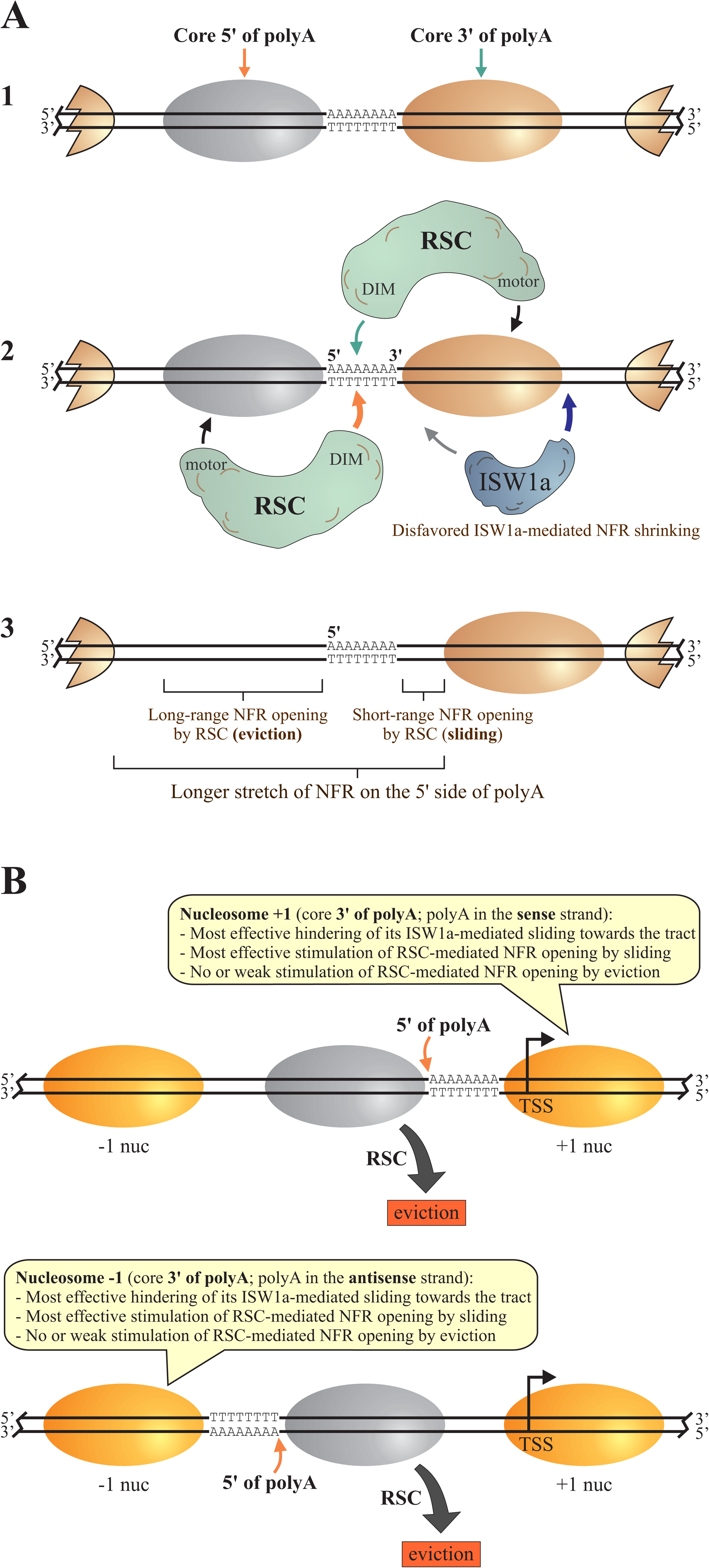
Proposed model for the effects of poly(dA:dT) tracts on ISW1a and RSC in the context of formation and maintenance of NFRs, and for the asymmetrical chromatin opening generated by these sequences. **(A)** General model proposed for poly(dA:dT) tracts present at gene promoters or any other genomic region. The model depicts three sequential steps for the effects of a single tract resulting in chromatin opening. 1) Upon histone deposition, DNA sequences passively locate poly(dA:dT) tracts preferentially in linker DNA regions. 2) Productive engagement of ISW1a on the poly(dA:dT)-containing linker is hindered by this sequence; the DNA-interaction module (DIM) of RSC preferentially binds to the tract-containing linker; for the core located 5’ of polyA, the tract mostly stimulates eviction activity of RSC; for the core located 3’ of polyA, the tract mostly stimulates sliding activity of RSC. 3) Eviction activity of RSC on the core located 5’ of polyA results in a long-range chromatin opening; RSC-mediated sliding activity on the core located 3’ of polyA results in a short-range opening; NFR shrinking by ISW1a is hindered by the tract. The histone octamer represented by the grey oval stands for the octamer evicted by RSC action. The fragments of nucleosomes shown in the right and left ends represent neighboring nucleosomes and do not correspond to any special nucleosome structure. **(B)** Tailored version of the model, focusing on the characteristic enrichment pattern of polyT and polyA present in the sense strand of gene promoters of *S. cerevisiae*. This tailored version of the proposed model highlights the mechanism that would underlie the enrichment of polyT upstream and polyA downstream the axis of gene promoter NFRs. The upper panel represents the effect of a tract on the +1 nucleosome and the nucleosome to be disassembled by RSC action (grey oval). The lower panel represents the effect of a tract on the-1 nucleosome and the nucleosome to be disassembled by RSC action (grey oval).

Regarding RSC and asymmetrical chromatin opening around poly(dA:dT) tracts, we propose the following mechanism, based in our findings and the known properties of the complex (Fig 8A). First, the DIM of RSC binds to a tract-containing linker, making this linker the exit DNA in the remodeling process. This binding ensures that this linker will be expanded, leading to NFR formation, independent of the adjacent core bound by the ATPase motor of the complex. Now, depending on the core bound by the motor, RSC will mainly expand this linker by sliding (if the core bound is 3’ of polyA) or by eviction of the histone octamer (if the core bound is 5’ of polyA). Chromatin opening would be of a short range in the case of sliding, as the stimulatory effect of the tract falls down as its distance to the core increase over 40 bp. On the other hand, eviction of the histone octamer clears a longer stretch of DNA. Thereafter, this differential effect on the nucleosome cores located besides the tract would lead to asymmetrical opening of this linker DNA region (Fig 8A). The same mechanism would explain why it is more common to find polyT upstream the midpoint of NFRs and polyA downstream this point at gene promoter NFRs, as well as low rates of RSC-mediated eviction of-1 and +1 nucleosomes (Fig 8B). Additionally, histone acetylation could also prevent eviction of-1 and +1 nucleosomes by RSC (Lorch *et al*, 2018). Future studies will be required to confirm whether the differential effects given by the two alternative tract orientations lead to differential biochemical outcomes of RSC *in vivo* (sliding versus eviction), in order to confirm whether this duality conforms the underlying mechanism of asymmetrical chromatin opening catalyzed by this complex around poly(dA:dT) tracts.

In the case of ISW1a, *in vivo* studies have found both, shrinking and widening of the NFR mediated by this complex (Tirosh *et al*, 2010; Yen *et al*, 2012; Parnell *et al*, 2015). Regarding these apparently contradictory findings, our remodeling assays and bioinformatics analyses support a view where ISW1a acts by shrinking tract-lacking NFRs and broadening tract-containing NFRs. Interestingly, in those genes that displayed upstream mobilization of the +1 nucleosome by ISW1a (NFR shrinking), there is a higher frequency of poly(dA:dT) tracts right downstream this nucleosome. This pattern suggests that there might be genes where poly(dA:dT) tracts, located downstream the +1 nucleosome, could assist ISW1a-mediated mobilization of this nucleosome towards the NFR, with the caveat that positioning of +1 nucleosomes obtained upon *in vitro* reconstitution of the *S. cerevisiae* genome does not necessarily match their *in vivo* positioning. Regarding the differential effect given by the two alternative tract orientations, the higher frequency of polyA in the sense strand closely upstream the +1 nucleosome and of polyT closely downstream the-1 nucleosome fits well with the stronger hindering of ISW1a given by the orientation “core 3’ of polyA” observed in our analyses (Fig 8B). In this context, recent studies have pointed that other CRCs play a more relevant role in the positioning of the +1 nucleosome, such as Ino80 and ISW2 (Kubik *et al*, 2019; Oberbeckmann *et al*, 2021a), making it interesting to study the effects of poly(dA:dT) tracts on nucleosome remodeling activity of these and other CRCs. Regarding this, it has been recently shown that a poly(dA:dT) tract placed near the border of a nucleosome core or on linker DNA exerts an inhibitory effect on Ino80 activity, although the effect of tract orientation was not tested (Kunert *et al*, 2022). Finally, it is worth to mention that other factors could also affect the action of ISW1a at gene promoter NFRs, such as transcription factors, including general regulatory factors (Li *et al*, 2015; Krietenstein *et al*, 2016).

Several relevant questions are raised by our findings. One is whether other DNA sequences exert effects on RSC in a way similar to this found for poly(dA:dT) tracts. Another question refers to the existence of two forms of the RSC complex, RSC1 and RSC2 (Schlichter *et al*, 2020). Our assays were performed using the RSC2 complex. Thus, it will be of interest to compare the effect of poly(dA:dT) sequences on the RSC1 and RSC2 complexes. An additional relevant question is whether poly(dA:dT) tracts and/or other DNA sequences exert similar effects on CRCs of higher eukaryotes. Interestingly, in the case of mammals, NFRs are also asymmetrically positioned around poly(dA:dT) tracts, but displaying a longer stretch of NFR on the 3’ side of polyA (de Boer & Hughes, 2014).

Taken together, our findings give insight into the mechanisms by which poly(dA:dT) tracts affect ATP-dependent nucleosome remodeling activity and also support the view that defined DNA sequences play roles in both passive and active processes that govern nucleosome positioning, which in turn affect transcriptional activity and other processes involving access to DNA. In addition, our findings suggest that defined DNA sequences are strategically located at gene promoters and other genomic regions, determining in which direction nucleosome sliding should proceed and even determining the biochemical outcome of the nucleosome remodeling process (sliding versus histone octamer ejection).

## Materials and Methods

### Protein complexes

The *S. cerevisiae* ISW1a and RSC complexes were obtained by tandem affinity purification from Ioc3-TAP and Rsc2-TAP strains, respectively (Table S2), as previously described (Rigaut *et al*, 1999). Purified complexes were analyzed as detailed in our previous report (Amigo *et al*, 2022). For both complexes, a portion of a purification was extensively concentrated (Microcon Ultracel YM-10, Amicon-Millipore), quantified by SDS– PAGE/Coomassie staining and then used as standard for Western blot quantification of the purified complex in that and further purifications.

### Plasmids, probes and nucleosome reconstitution

DNA probes of different lengths and harboring the 601 nucleosome positioning sequence located at different positions were generated by PCR using distinct plasmids as templates. The 147 bp positioning region of the 601 sequence was defined as previously described (Li & Widom, 2004). The plasmids and primer sets used are depicted in Table S1. Before PCR amplification, one of the primers used in each reaction was labeled on its 5’ end using [γ-^32^P]-ATP (Perkin-Elmer NEG035C or ARC ARP0102B). Nucleosome reconstitution was carried out by the octamer transfer method, as previously described (Gutierrez *et al*, 2007). Oligonucleosomes used as histone donors for reconstitution were obtained from HeLa cells as described elsewhere (Utley *et al*, 1996). All the reconstitution reactions used for this study were carried out using 0.5 pmol of probe and 1.5 µg of oligonucleosomes. Once reconstituted, the nucleosome probes (and mock-reconstituted probes) contain 4 fmol/μL of probe and 12 ng/μL of non-labeled oligonucleosomes (the latter value in terms of DNA content).

### Competitive nucleosome reconstitution

Nucleosome reconstitution for these assays was performed as stated above, with the following changes: Donor oligonucleosomes were obtained from chicken erythrocytes as previously described (Gutierrez *et al*, 2000). The amount of oligonucleosomes for the reconstitution reaction was set as the amount required to obtain 30% reconstitution of the 147 bp 5s probe. This probe corresponds to the dominant positioning region of the *Lytechinus variegatus* 5s rDNA, when reconstituted into mononucleosomes (Simpson & Stafford, 1983; Hansen *et al*, 1989). All the different probes were reconstituted simultaneously in each independent assay. Three independent assays were performed (two in triplicate, one in duplicate). In each assay, the components used for reconstitution were taken from the same master mix. Relative equilibrium constant values (relative Keq) and relative free energies (ΔΔG°) were calculated as described by Thastrom and co-workers (Thastrom *et al*, 2004).

### Binding assays

In each binding reaction, a mix containing 6.9 μL of Remodeling buffer, 0.6 μL of ATP (Roche, 11140965001) or Adenosine-5’-O-(3-thiotriphosphate) (Roche, 11162306001), 1.5 μL of TF buffer, 0.5 μL of CADS buffer, 3 μL of CRC or CRC buffer and 2.5 μL of probe was incubated for 45 min at 30°C (see the composition of each solution used in the mixes in the Supplementary Information). Final concentration of the different components of this mixture was: 10.4 mM HEPES-KOH (pH 7.4–7.9), 3.9 mM Tris-Cl (pH 7.4–8.0), 83.6 mM NaCl, 36.4 mM KCl, 5.1 mM MgCl_2_, 2 mM ATP (or ATP-γ-S), 0.2 mM Mg(CH_3_COO)_2_, 0.05% NP-40, 10% glycerol, 1 μM ZnCl_2_, 0.2 mM EDTA (pH 8.0), 0.4 mM EGTA, 73 μg/mL BSA, 1.2 mM imidazole, 2.1 mM DTT, and 0.45 mM PMSF. The samples were then subjected to electrophoresis in non-denaturing polyacrylamide gel (200V, 0.3x TBE, 3.5% acrylamide, 60:1 AA:Bis proportion) in cold room. Afterwards, the gel was dried and autoradiographed on film.

### Nucleosome remodeling assays

For remodeling reactions, a mix containing the same composition described for binding assays was used. Each reaction was incubated for 45 min at 30°C, except for assays where the time is specifically indicated in a figure. After this incubation, a mix (0.7 μL) containing 750 ng salmon sperm DNA and 500 ng long oligonucleosomes was added, incubating for 20 additional minutes at 30 °C. The samples were then subjected to electrophoresis in non-denaturing polyacrylamide gel (200V, 0.3x TBE, 5% acrylamide, 40:1 AA:Bis proportion) in cold room. Afterwards, the gel was dried and then scanned using phosphor screen and Molecular Imager FX (BioRad). Densitometric analyses were performed using Quantity One software (BioRad). Film autoradiography was also performed. The extent of nucleosome sliding was calculated as the ratio of the signal of slid mononucleosome band over the signal of all mononucleosome bands in the lane. The extent of nucleosome eviction was calculated by subtracting the free DNA signal of the probe present in the absence of RSC from the free DNA signal obtained in the presence of this complex. The resulting value was divided by the nucleosomal DNA signal present in the absence of complex, obtaining the fraction of nucleosomal DNA that is converted to free DNA. Values of sliding or eviction extent were made relative to the average of all reactions present within an individual assay before combining data with the other independent assays. In time-course assays, after the remodeling and removing steps, the mixes were placed on ice until the end of all reactions. Afterwards, the samples were subjected to gel electrophoresis and subsequent analysis as described above.

### Determination of nucleosome sliding directionality

Nucleosome remodeling reactions were performed as described above. Once removing reactions were done, 10 U of a specific restriction enzyme were added and an additional incubation was performed for 30 min at 30°C. Afterwards, the samples were subjected to gel electrophoresis and subsequent analysis as described for remodeling assays.

### Bioinformatics analyses

For determination of nucleosome positioning and its change in the presence of ISW1a, datasets obtained from a study of Krietenstein and co-workers (GEO dataset GSE72106) were used (Krietenstein *et al*, 2016). The datasets used were MNase-ChIP-H3-seq on *S. cerevisiae* genome reconstituted into nucleosomes (SGD), replicate 1, replicate 2, and replicate 3 (Sequence Read Archive accession number SRR2164369, SRR2164396, and SRR2164397, respectively) and MNase-seq on reconstituted genome in the presence of purified ISW1a, replicate 1 and replicate 2 (Sequence Read Archive accession number SRR2164386 and SRR2164392, respectively). These datasets were processed as detailed in the Supplementary Information. Mitochondrial genome sequences and ribosomal genes were excluded from the analysis. To obtain nucleosome positions, the nf-core/mnaseseq pipeline from the NextFlow hub was executed through Docker. The +1 nucleosome was defined as the nucleosome dyad peak that was closest to a TSS in a window of-80 bp to +140 bp. The-1 nucleosome was defined as the upstream nucleosome dyad peak that was closest to the +1 nucleosome. All conditions were processed using these criteria. Changes in the translational location of nucleosomes were defined as the difference in nucleosome position between the absence (SGD) and the presence of purified ISW1a for +1 and-1 nucleosomes. The obtained changes in nucleosome positioning were classified in two groups: more than 20 bp upstream and more than 20 bp downstream the original position (SGD), for both +1 and-1 nucleosomes. For the analysis of poly(dA:dT) tracts frequency, windows of 250 bp around nucleosome dyad peaks on SGD were defined and the corresponding sequences were obtained in FASTA format (500 bp window). PolyA and polyT tracts were defined to be at least 6 nucleotides (5’-AAAAAA-3’ and 5’-TTTTTT-3’). Sequences around nucleosome dyad were binned in 25 bp intervals and polyA or polyT tract frequency was plotted for each bin. See more details for each step in the Supplementary Information.

## Acknowledgements

This work was supported by grants National Agency for Research and Development (ANID) [FONDECYT Regular 1180911 to J.L.G.], ANID [FONDECYT Iniciación 11190401 to E.T.], ANID [Subvención a la Instalación en la Academia SA77210106 to C.F.], Universidad de Concepción [VRID-Enlace 216.037.020-1.0 to J.L.G.], and ANID [Scholarship for Ph.D. students 21170786 to R.A. and 21230435 to F.R.].

## Data availability

This study includes no data deposited in external repositories.

## Supplementary information

Supplementary information is available in a separate file.

## Disclosure and competing interests statement

The authors declare that they have no conflict of interest.

## Notes

### Competing Interest Statement

The authors have declared no competing interest.

