## Supplementary Information for "Poly(dA:dT) tracts differentially modulate nucleosome remodeling activity of RSC and ISW1a complexes, exerting tract orientation-dependent and -independent effects"

#### Materials and Methods

##### Composition of buffers mixed in binding (EMSA) and nucleosome remodeling assays

(final concentration of each component is given in the main text).

Remodeling buffer (6.9 µL): 20 mM HEPES-KOH (pH 7.9), 57.3 mM KCl, 0.5 mM PMSF, 2 mM DTT, 0.05 % NP-40, 8.7 % Glycerol, 11 mM $MgCl_2$, 100 µg/mL BSA.

His-protein or TF buffer (1.5 µL): 10 mM HEPES-KOH (pH 7.4), 100 mM KCl, 20 % Glycerol, 10 µM $ZnCl_2$, 10 mM Imidazole, 100 µg/mL BSA, 1 mM DTT, 0.2 mM PMSF.

Buffer CADS (0.5 µL): 6.96 mM HEPES-KOH (pH 7.5), 6.52 mM Tris-Cl (pH 7.4), 208.7 mM NaCl, 0.5 mM PMSF, 3.61 mM DTT, 1 mM EDTA (pH 8.0), 0.03 % NP-40, 10 % Glycerol.

Chromatin remodeling complex or CRC buffer (3 µL): 10 mM Tris-Cl (pH 8.0), 300 mM NaCl, 1 mM $Mg(CH_3COO)_2$, 1 mM Imidazole, 2 mM EGTA, 0.1 % NP-40, 0.5 mM PMSF, 0.5 mM DTT, 10 % Glycerol.

Deionized water, ATP or ATP-γ-S (0.6 µL): 50 mM ATP (or ATP-γ-S).

Probe (2.5 µL): 10 mM Tris-Cl (pH 7.4), 1 mM EDTA, 0,5 mM PMSF, 5 mM DTT, 0,05% NP-40, 10 % Glycerol, 100 µg/mL BSA, 100 mM NaCl.

Total reaction volume: 15 µL.

##### Bioinformatics analyses

For the determination of nucleosome positioning and its change in the presence of ISW1a, datasets obtained from Krietenstein *et al*, 2016 (GEO dataset GSE72106) were used. We first downloaded fastq datasets using the commands prefetch and fastq-dump belonging to SRA tools (https://github.com/ncbi/sra-tools). For SGD condition, MNase-ChIP-H3-seq of *S. cerevisiae* genomic DNA reconstitution, replicate 1, replicate 2, and replicate 3 (Sequence Read Archive accession number SRR2164369 [PSU43201], SRR2164396 [PSU39301], and SRR2164397 [PSU52904] respectively) were used. In addition, MNase-seq of reconstituted genomic DNA in the presence of purified ISW1a, replicate 1 and replicate 2 (Sequence Read Archive accession number SRR2164386 [PSU51101] and SRR2164392 [PSU51302], respectively) were used.

Next, we trimmed our fastq datasets using the fastp tool with default parameters (1). After fastq QC and trimming, we downloaded the sacCer3 genome and GTF annotation file from the UCSC database. To exclude the mitochondrial genome sequences from the genome assembly, we converted the FASTA genome assembly into a tabular format using seqkit (2); we used the grep command with the --invert-match -F "chrM" flag and regenerated the FASTA file from the tabular file. We also removed genomic positions corresponding to the rRNA genes (Chromosome XII: 451535-489509) from the alignments by building a bed file containing the referred region (ribosomal_DNA.bed), which served as a blacklisted region in the genomic alignments.

After these steps, we installed and ran the automated nf-core/mnaseseq pipeline from the NextFlow hub through Docker (https://nf-co.re/mnaseseq) (3). We downloaded the design_se.csv file from the nf-core/mnaseseq repository on GitHub (https://github.com/nf-core/mnaseseq), which contained the group and replicate information to input the processed fastq files. With this template, we created a file manifest (design.csv file) for all replicates, comparing SGD versus ISW1a. We then ran the mnaseseq pipeline using the design file and specified parameters such as --single_end yes, --fasta sacCer3_non_chrM.fa, --gtf sacCer3_ncbiRefSeq_non_chrM.gtf, --blacklist ribosomal_DNA.bed, and --skip_trimming yes. The nf-core/mnaseseq pipeline included as sequential steps: quality control of raw reads (FastQC: https://www.bioinformatics.babraham.ac.uk/projects/fastqc/); adapter trimming (Trim Galore!: https://github.com/FelixKrueger/TrimGalore); alignment to a reference genome (BWA) (4); marking and removing duplicates (picard tools: http://broadinstitute.github.io/picard/); reads filtering, including removal of reads that map to blacklisted regions (ribosomal_DNA.bed), duplicates, unmapped, and reads that contain too many mismatches (SAMtools, BAMTools, Pysam) (5,6); alignment-level quality control and library complexity estimation (picard, Preseq) (7); and finally calling nucleosome positions and generating occupancy profile plots (DANPOS2) (8). To obtain nucleosome positions, the pipeline also merged replicates, which was the final output used in this work.

Once nucleosome positions were obtained, all conditions were processed as follows: For each gene, the +1 nucleosome was defined as the nucleosome dyad peak that was closest to the TSS in a window of -80 bp to +140 bp and -1 nucleosome was defined as the upstream nucleosome dyad peak that was closest to the +1 nucleosome. Minus 1 nucleosomes at a distance greater than 1000 bp were manually discarded. SGD and ISW1a datasets were aligned, and we stayed with genes present in both datasets (4064 genes). Movement of nucleosomes was defined as the difference in nucleosome position between the absence and the presence of purified ISW1a for +1 and, separately, -1 nucleosome. Changes in nucleosome positioning were classified in two groups, separately for +1 and -1 nucleosomes: more than 20 bp upstream and more than 20 bp downstream.

For the analysis of poly(dA:dT) tracts frequency, first Fasta sequence of genes in a window of -250 to +250 bp around the dyad peaks were obtained using bedtools getfasta; -s option was used to obtain the reverse complementary of minus strand genes. fasta-grep (MEME suite 5.5.0) was used to find poly(dA:dT) tracts on the sense strand; -dna, -p, and -norc options were used. PolyT and polyA were defined to be at least 6 nucleotides: 5’-TTTTTT-3’ and 5’-AAAAAA-3’, respectively. The analysis was performed for six clusters of genes: 1) SGD (-1 nucleosome; 4064 genes), 2) SGD (+1 nucleosome; 4064 genes), 3) ISW1a upstream movement (-1 nucleosome; 1793 genes), 4) ISW1a upstream movement (+1 nucleosome; 890 genes), 5) ISW1a downstream movement (-1 nucleosome; 854 genes), 6) ISW1a downstream movement (+1 nucleosome; 1543 genes). Sequences around nucleosome dyad were binned in 25 bp (20 windows for 500 nucleotides) intervals and polyA or polyT frequencies were plotted.

#### Figure legends

**Figure S1. Description of the nomenclature used in this work to denote locations and orientations of poly(dA:dT) tracts relative to nucleosomes and defined genomic regions. (A)** Dominant positioning region of the Widom's 601 sequence (147 bp). The upper strand is defined as that starting with the sequence ACAG. **(B)** Schematic representation depicting equivalent denominations for each location and orientation of poly(dA:dT) tracts. The ellipse represents the translational position of the nucleosome core. The gray line represents the 601 nucleosome positioning sequence. SHL = superhelix location.

**Figure S2. Poly(dA:dT) tracts located within the nucleosome core (SHL 5.5 proximal to linker DNA) exhibit an overall weaker effect on the activity of ISW1a and RSC complexes, as compared to their presence on linker DNA. (A)** Schematic representation of the nucleosome probes used in the assays. **(B, C)** Nucleosome remodeling assays visualized by electrophoresis in non-denaturing polyacrylamide gels, performed for ISW1a (B) and RSC (C). The probe used in each reaction is depicted at the top of each gel picture, where absence or presence of the corresponding remodeling complex is also depicted (1.5 nM ISW1a, 2 nM RSC). Each image is representative of three independent assays, performed under the same conditions. Migrations of alternative forms of the nucleosome probe, which correspond to different translational positions of the histone octamer, are indicated schematically at the right of each picture. The graph at the bottom of each gel picture depicts quantification of sliding extent from the corresponding assay (see Fig 1B for information regarding quantification strategy). Bars in the graphs display the average of three independent assays for each condition analyzed (n = 3). Error bars represent one standard deviation. Asterisks denote statistically significant differences (***p<0.001) and n.s. statistically non-significant difference between the indicated conditions, as deducted from ANOVA with Tukey's multiple comparisons test.

**Figure S3. The effect of a poly(dA:dT) tract on the sliding activity of ISW1a and RSC has a tract length-dependency. (A)** Schematic representation of the nucleosome probes used in (B) and (C). **(B, C)** Nucleosome remodeling assays visualized by electrophoresis in non-denaturing polyacrylamide gels, testing the effect of poly(dA:dT) tracts on ISW1a (B) and RSC (C) sliding activity, comparing 15 bp to 7 bp tract length. The probe used in each reaction is depicted at the top of each gel picture, where absence or presence of the corresponding remodeling complex is also depicted (1.5 nM ISW1a, 2 nM RSC). Each image is representative of three independent assays, performed under the same conditions. Migrations of alternative forms of the nucleosome probe, which correspond to different translational positions of the histone octamer, are indicated schematically at the right of each picture. The graph below each gel picture depicts quantification of sliding extent from the corresponding assay (see Fig 1B for information regarding quantification strategy). Bars in the graphs display the average of three independent assays for each condition analyzed (n = 3). Error bars represent one standard deviation. Asterisks denote statistically significant differences (*p<0.05; **p<0.01; ***p<0.001) and n.s. statistically non-significant difference between the indicated conditions, as deducted from ANOVA with Tukey's multiple comparisons test.

**Figure S4. RSC displays the same affinity to the two alternative poly(dA:dT) tract orientations on naked DNA probes.** Binding analysis (EMSA) for RSC using naked DNA probes (mock reconstitutions). The probes used in this assay are the same used in the binding assay shown in Fig 3B, but probes are at the form of mononucleosomes in that assay. A schematic representation of these probes can be seen in the left panel of Fig 1A. The probe used in each reaction is depicted at the top of the gel picture, where absence or presence of ATP, ATP- γ-S and the remodeling complex (8 nM) is also depicted. The gel image is representative of three independent assays, performed under the same conditions. Migration of unbound probe (DNA) and the RSC/DNA complex are indicated at the right of the gel picture.

**Figure S5. RSC action on nucleosomal probes harboring the orientation corresponding to "core 5' of polyA" results in higher generation of smear, as compared to the opposite tract orientation. (A)** Schematic representation of the nucleosome probes used in the assays. **(B, C)** Nucleosome remodeling assays visualized by electrophoresis in non-denaturing polyacrylamide gels, testing the effect of poly(dA:dT) tracts on the activity of ISW1a (B) and RSC (C) complexes. The probe used in each reaction is depicted at the top of each gel picture, where absence or presence of the corresponding remodeling complex is also depicted (1.5 nM ISW1a, 2 nM RSC). Each image is representative of at least three independent assays, performed under the same conditions. Migrations of alternative forms of the nucleosome probe, which correspond to different translational positions of the histone octamer, are indicated schematically at the right of the picture. 2x: Remodeling complex concentration two times the concentration used in the other reactions. The gel pictures correspond to image acquisition using phosphor imager. The graph at the right side of each gel image corresponds to a densitometric profile of selected lanes. The column graphs in (C) correspond to quantification of sliding extent, carried out using the regular strategy (top, see Fig 1B for information regarding quantification strategy) or a strategy considering the smear generated between the signal of the nucleosomal probe and the loading well (bottom). Bars in the graphs display the average of five replicates for each condition analyzed. Error bars represent one standard deviation. Asterisks denote statistically significant differences (**$p<0.01$; ***$p<0.001$; ****$p<0.0001$), as deducted from ANOVA with Tukey's multiple comparisons test.

**Figure S6. The differential stimulatory effect given by the two alternative tract orientations on RSC sliding activity is maintained in mononucleosome probes harboring linker DNA on both sides of the core. (A)** Schematic representation of the nucleosome probes used in the assay. **(B)** Nucleosome remodeling assay visualized by electrophoresis in a non-denaturing polyacrylamide gel, testing the effect of a poly(dA:dT) tract, located in the downstream linker, on RSC sliding activity. The probe used in each reaction is depicted at the top of the gel picture, where absence or presence of the remodeling complex (2 nM) is also depicted. The image is representative of three independent assays, performed under the same conditions. Migrations of alternative forms of the nucleosome probe, which correspond to different translational positions of the histone octamer, are indicated schematically at the right of the picture; the directionality of histone octamer mobilization is indicated with base in the analysis shown in Figs 4C and 4E. **(C)** Quantification of sliding extent (see Fig 1B for information regarding quantification strategy). Bars in the graph display the average of three independent assays for each condition analyzed ($n = 3$). Error bars represent one standard deviation. Asterisks denote statistically significant differences (*$p<0.05$; ****$p<0.0001$), as deducted from ANOVA with Tukey's multiple comparisons test.

**Figure S7. RSC also slides the histone octamer away from a poly(dA:dT) tract on laterally positioned nucleosome probes. (A)** Schematic representation of the nucleosome probes used in the assays. **(B)** Schematic representation depicting changes in accessibility to restriction enzymes depending on upstream or downstream mobilization of the histone octamer. Names of restriction enzymes presented in gray represent absence of accessibility, while names presented in black represent access of the corresponding restriction site. The numbers below the names of restriction enzymes indicate the size of labeled DNA segments resulting from digestion by the corresponding enzyme, relative to the 5' end of the upper strand (labeled DNA end, represented by an asterisk). Note that cutting of the nucleosomal probe by *Pml*I would result in a labeled fragment corresponding to a very short naked DNA fragment (22 bp, which runs out the gel). On the other hand, cutting of the nucleosomal probe by *Xho*I results in a 163 bp labeled fragment corresponding to a nucleosome. **(C, D)** Nucleosome remodeling assays testing directionality of RSC sliding activity by changes in accessibility to restriction enzymes, visualized by electrophoresis in non-denaturing polyacrylamide gels. The probe used in each assay is depicted at the top of each gel picture, where the presence of RSC (2 nM) and/or a defined restriction enzyme is also depicted. Each image is representative of three independent assays, performed under the same conditions. Migration of different species is depicted at the right of each gel picture. In both assays, lanes 1 to 3 correspond to probe at the form of naked DNA (mock reconstitution).

**Figure S8. Poly(dA:dT) tracts do not hinder ISW1a activity when located in the linker that becomes exit DNA during the sliding process. (A)** Schematic representation of the nucleosome probes used in the assay. **(B)** Nucleosome remodeling assay visualized by electrophoresis in a non-denaturing polyacrylamide gel, testing the effect of a poly(dA:dT) tract, located in the shorter linker upstream the core, on ISW1a sliding activity. The probe used in each reaction is depicted at the top of the gel picture, where absence or presence of the remodeling complex (1.5 nM) is also depicted. The image is representative of three independent assays, performed under the same conditions. Migrations of alternative forms of the nucleosome probe, which correspond to different translational positions of the histone octamer, are indicated schematically at the right of the picture. **(C)** Quantification of sliding extent (see Fig 1B for information regarding quantification strategy). Bars in the graph display the average of three independent assays for each condition analyzed ($n = 3$). Error bars represent one standard deviation. Asterisks denote statistically significant differences (***$p<0.001$), as deducted from ANOVA with Tukey's multiple comparisons test.

**Figure S9. In a nucleosome probe harboring tracts flanking the core in the orientation "core 3' of polyA", RSC action results in a pattern that lacks the fastest migrating band.** These assays are complementary to those shown in figures 6A and 6B, where both tract orientations are compared. Here, a probe harboring two tracts, one in each linker, is compared to probes harboring no tract or harboring a tract in only one of the linkers, always in the same orientation (core 3' of polyA). Taken together, the analyses show that, although the presence of two tracts with this orientation exerts a stimulatory effect that is stronger than the presence of only one tract, there is no generation of the fastest-migrating remodeled state in this probe harboring two tracts surrounding the core. **(A)** Schematic representation of the nucleosome probes used in the assays. **(B)** *Left panel*: Nucleosome remodeling assay visualized by electrophoresis in a non-denaturing polyacrylamide gel. The probe used in each reaction is depicted at the top of the gel picture, where absence or presence of the remodeling complex (2 nM) is also depicted. The image is representative of three independent assays, performed under the same conditions. Migrations of alternative forms of the nucleosome probe, which correspond to different translational positions of the histone octamer, are indicated schematically at the right of the picture. *Right panel*: Quantification of sliding extent (see Fig 1B for information regarding quantification strategy). Bars in the graph display the average of three independent assays for each condition analyzed ($n = 3$). Error bars represent one standard deviation. Asterisks denote statistically significant differences (**$p<0.01$; ****$p<0.0001$), as deducted from ANOVA with Tukey's multiple comparisons test. **(C)** Time-course analysis. See legend in (B) for a general description of the remodeling assay and data analysis. **(D)** Additional time-course analysis, including the control probe harboring no tracts and performing longer incubation times. The long incubation times were intended to show that RSC action on the control probe does result in the fastest migrating remodeling state, despite the very weak activity of RSC on this probe, further confirming that this remodeled state is not obtained in the case of the probe harboring tract on both sides of the core (both tracts in the orientation "core 3' of polyA"). See legend in (B) for a general description of the remodeling assay.

**Table S1. Sequence information of primers and template plasmid used for generation of each probe.**

| Probe name | Plasmid name and sequence harboring the region amplified in PCR reaction | Primers |
|---|---|---|
| 0-NC-70 | pGEM-3Z/601-Gal4<br>5’ggtcgctgttcaatacatgcacaggatgtatatatctgacacgtgcctggagactagggagtaatccc<br>cttggcggttaaaacgcgggggacagcgcgtacgtgcgtttaagcggtgctagagctgtctacgaccaat<br>tgagcggcctcggcaccgggattctccagggcggccgcgtatctcgagcatcggaggacagtcctccgc<br>ggacctgcaggcatgcaagcttgagtattctatagtgtcacctaaatagc | Forward:<br>5’ACAGGATGTATATATCTGA<br>CACGTGCCTGG<br>Reverse:<br>5’GAATACTCAAGCTTGCATG<br>CCTG |
| 0-NC-70$A_{15}$ | pGEM-3Z/601-Gal4-A15-2in<br>5’ggtcgctgttcaatacatgcacaggatgtatatatctgacacgtgcctggagactagggagtaatccc<br>cttggcggttaaaacgcgggggacagcgcgtacgtgcgtttaagcggtgctagagctgtctacgaccaat<br>tgagcggcctcggcaccgggattctccaaaaaaaaaaaaaaactcgagcatcggaggacagtcctccg<br>cggacctgcaggcatgcaagcttgagtattctatagtgtcacctaaatagc | Forward:<br>5’ACAGGATGTATATATCTGA<br>CACGTGCCTGG<br>Reverse:<br>5’GAATACTCAAGCTTGCATG<br>CCTG |
| 0-NC-70$T_{15}$ | pGEM-3Z/601-Gal4-T15-2in<br>5’ggtcgctgttcaatacatgcacaggatgtatatatctgacacgtgcctggagactagggagtaatccc<br>cttggcggttaaaacgcgggggacagcgcgtacgtgcgtttaagcggtgctagagctgtctacgaccaat<br>tgagcggcctcggcaccgggattctccttttttttttttttctcgagcatcggaggacagtcctccgcggac<br>ctgcaggcatgcaagcttgagtattctatagtgtcacctaaatagc | Forward:<br>5’ACAGGATGTATATATCTGA<br>CACGTGCCTGG<br>Reverse:<br>5’GAATACTCAAGCTTGCATG<br>CCTG |
| 0-NC-70$T_{10}$ | pGEM-3Z/601-Gal4-T10-2in<br>5’ggtcgctgttcaatacatgcacaggatgtatatatctgacacgtgcctggagactagggagtaatccc<br>cttggcggttaaaacgcgggggacagcgcgtacgtgcgtttaagcggtgctagagctgtctacgaccaat<br>tgagcggcctcggcaccgggattctcctttttttttcgtatctcgagcatcggaggacagtcctccgcgga<br>cctgcaggcatgcaagcttgagtattctatagtgtcacctaaatagc | Forward:<br>5’ACAGGATGTATATATCTGA<br>CACGTGCCTGG<br>Reverse:<br>5’GAATACTCAAGCTTGCATG<br>CCTG |
| 0-NC-70$T_7$ | pGEM-3Z/601-Gal4-T7-2in<br>5’ggtcgctgttcaatacatgcacaggatgtatatatctgacacgtgcctggagactagggagtaatccc<br>cttggcggttaaaacgcgggggacagcgcgtacgtgcgtttaagcggtgctagagctgtctacgaccaat<br>tgagcggcctcggcaccgggattctccttttttccgcgtatctcgagcatcggaggacagtcctccgcgg<br>acctgcaggcatgcaagcttgagtattctatagtgtcacctaaatagc | Forward:<br>5’ACAGGATGTATATATCTGA<br>CACGTGCCTGG<br>Reverse:<br>5’GAATACTCAAGCTTGCATG<br>CCTG |
| 0-NC-70$T_5$ | pGEM-3Z/601-Gal4-T5-2in<br>5’ggtcgctgttcaatacatgcacaggatgtatatatctgacacgtgcctggagactagggagtaatccc<br>cttggcggttaaaacgcgggggacagcgcgtacgtgcgtttaagcggtgctagagctgtctacgaccaat<br>tgagcggcctcggcaccgggattctcctttttggccgcgtatctcgagcatcggaggacagtcctccgcgg<br>acctgcaggcatgcaagcttgagtattctatagtgtcacctaaatagc | Forward:<br>5’ACAGGATGTATATATCTGA<br>CACGTGCCTGG<br>Reverse:<br>5’GAATACTCAAGCTTGCATG<br>CCTG |
| 0-NC-19-$T_{15}$ | pGEM-3Z/601-T15-19<br>5’ggtcgctgttcaatacatgcacaggatgtatatatctgacacgtgcctggagactagggagtaatccc<br>cttggcggttaaaacgcgggggacagcgcgtacgtgcgtttaagcggtgctagagctgtctacgaccaat<br>tgagcggcctcggcaccgggattctccagggcggccgcgtatctcgagtttttttttttttttctccgcggac<br>ctgcaggcatgcaagcttgagtattctatagtgtcacctaaatagc | Forward:<br>5’ACAGGATGTATATATCTGA<br>CACGTGCCTGG<br>Reverse:<br>5’GAATACTCAAGCTTGCATG<br>CCTG |
| 0-NC-40-$T_{15}$ | pGEM-3Z/601-Gal4-T15-40<br>5’ggtcgctgttcaatacatgcacaggatgtatatatctgacacgtgcctggagactagggagtaatccc<br>cttggcggttaaaacgcgggggacagcgcgtacgtgcgtttaagcggtgctagagctgtctacgaccaat<br>tgagcggcctcggcaccgggattctccagggcggccgcgtatctcgagcatcggaggacagtcctccgct<br>tttttttttttttcaagcttgagtattctatagtgtcacctaaatagc | Forward:<br>5’ACAGGATGTATATATCTGA<br>CACGTGCCTGG<br>Reverse:<br>5’GAATACTCAAGCTTGAAA |

| | | |
|---|---|---|
| 0-NC-80 | pGEM-3Z/601-Gal4:<br>5’ggtcgctgttcaatacatgcacaggatgtatatatctgacacgtgcctggagactagggagtaatccc<br>cttggcggttaaaacgcgggggacagcgcgtacgtgcgtttaagcggtgctagagctgtctacgaccaat<br>tgagcggcctcggcaccgggattctccagggcggccgcgtatctcgagcatcggaggacagtcctccgc<br>ggacctgcaggcatgcaagcttgagtattctatagtgtcacctaaatagcttggcgtaat | Forward:<br>5’ACAGGATGTATATATCTGA<br>CACGTGCCTGG<br>Reverse:<br>5’TGACACTATAGAATACTCA<br>AGC |
| 0-NCA$_{15}$-SHL5.5-80 | pGEM-3Z/601-A15-SHL5<br>5’ggtcgctgttcaatacatgcacaggatgtatatatctgacacgtgcctggagactagggagtaatccc<br>cttggcggttaaaacgcgggggacagcgcgtacgtgcgtttaagcggtgctagagctgtctacgaccaat<br>tgaaaaaaaaaaaaaaagggattctccagggcggccgcgtatctcgagcatcggaggacagtcctccg<br>cggacctgcaggcatgcaagcttgagtattctatagtgtcacctaaatagcttggcgtaat | Forward:<br>5’ACAGGATGTATATATCTGA<br>CACGTGCCTGG<br>Reverse:<br>5’TGACACTATAGAATACTCA<br>AGC |
| 0-NCT$_{15}$-SHL5.5-80 | pGEM-3Z/601-T15-SHL5<br>5’ggtcgctgttcaatacatgcacaggatgtatatatctgacacgtgcctggagactagggagtaatccc<br>cttggcggttaaaacgcgggggacagcgcgtacgtgcgtttaagcggtgctagagctgtctacgaccaat<br>tgtttttttttttttttgggattctccagggcggccgcgtatctcgagcatcggaggacagtcctccgcggac<br>ctgcaggcatgcaagcttgagtattctatagtgtcacctaaatagcttggcgtaat | Forward:<br>5’ACAGGATGTATATATCTGA<br>CACGTGCCTGG<br>Reverse:<br>5’TGACACTATAGAATACTCA<br>AGC |
| 70-NC-0 | pGEM-3Z/BNB-601<br>5’tgtaatacgactcactatagggcgaattcgagctcggtacccggggatcctatccgactggcacgcta<br>gcggtcgccgttcaatagatctaccggatgtatatatctgacacgtgcctggagactagggagtaatccc<br>cttggcggttaaaacgcgggggacagcgcgtacgtgcgtttaagcggtgctagagctgtctacgaccaat<br>tgagcggcctcggcaccgggattctccagggcggccgcgtatctcgagcatcggaggacagtcctccgc<br>ggacctgcaggcatgcaag | Forward:<br>5’GGCGAATTCGAGCTCGGT<br>Reverse:<br>5’CTGGAGAATCCCGGTGCC<br>GAG |
| 70A$_{15}$-NC-0 | pGEM-3Z/BNB-A15-601<br>5’tgtaatacgactcactatagggcgaattcgagctcggtacccggggatcctatccgactggcacgcta<br>gcggtcgccaaaaaaaaaaaaaaacggatgtatatatctgacacgtgcctggagactagggagtaatc<br>cccttggcggttaaaacgcgggggacagcgcgtacgtgcgtttaagcggtgctagagctgtctacgacc<br>aattgagcggcctcggcaccgggattctccagggcggccgcgtatctcgagcatcggaggacagtcctc<br>cgcggacctgcaggcatgcaag | Forward:<br>5’GGCGAATTCGAGCTCGGT<br>Reverse:<br>5’CTGGAGAATCCCGGTGCC<br>GAG |
| 70T$_{15}$-NC-0 | pGEM-3Z/BNB-T15-601<br>5’tgtaatacgactcactatagggcgaattcgagctcggtacccggggatcctatccgactggcacgcta<br>gcggtcgcctttttttttttttttcggatgtatatatctgacacgtgcctggagactagggagtaatccccttg<br>gcggttaaaacgcgggggacagcgcgtacgtgcgtttaagcggtgctagagctgtctacgaccaattga<br>gcggcctcggcaccgggattctccagggcggccgcgtatctcgagcatcggaggacagtcctccgcgga<br>cctgcaggcatgcaag | Forward:<br>5’GGCGAATTCGAGCTCGGT<br>Reverse:<br>5’CTGGAGAATCCCGGTGCC<br>GAG |
| 35-NC-35 | pGEM-3Z/BNB-601<br>5’gctcggtacccggggatcctatccgactggcacgctagcggtcgccgttcaatagatctaccggatgt<br>atatatctgacacgtgcctggagactagggagtaatccccttggcggttaaaacgcgggggacagcgcg<br>tacgtgcgtttaagcggtgctagagctgtctacgaccaattgagcggcctcggcaccgggattctccagg<br>gcggccgcgtatctcgagcatcggaggacagtcctccgcggacctgcaggcatgcaagc | Forward:<br>5’GACTGGCACGCTAGCGG<br>Reverse:<br>5’GGACTGTCCTCCGATGC |
| 35A$_{15}$-NC-35 | pGEM-3Z/BNB-A15-601<br>5’gctcggtacccggggatcctatccgactggcacgctagcggtcgccaaaaaaaaaaaaaaacggat<br>gtatatatctgacacgtgcctggagactagggagtaatccccttggcggttaaaacgcgggggacagcg<br>cgtacgtgcgtttaagcggtgctagagctgtctacgaccaattgagcggcctcggcaccgggattctcca<br>gggcggccgcgtatctcgagcatcggaggacagtcctccgcggacctgcaggcatgcaagc | Forward:<br>5’GACTGGCACGCTAGCGG<br>Reverse:<br>5’GGACTGTCCTCCGATGC |
| 35T$_{15}$-NC-35 | pGEM-3Z/BNB-T15-601<br>5’gctcggtacccggggatcctatccgactggcacgctagcggtcgccttttttttttttttttcggatgtatat<br>atctgacacgtgcctggagactagggagtaatccccttggcggttaaaacgcgggggacagcgcgtacg<br>tgcgtttaagcggtgctagagctgtctacgaccaattgagcggcctcggcaccgggattctccagggcgg<br>ccgcgtatctcgagcatcggaggacagtcctccgcggacctgcaggcatgcaagc | Forward:<br>5’GACTGGCACGCTAGCGG<br>Reverse:<br>5’GGACTGTCCTCCGATGC |
| 35-NC-35A$_{15}$ | pGEM-3Z/BNB-601-A15<br>5’gctcggtacccggggatcctatccgactggcacgctagcggtcgccgttcaatagatctaccggatgt<br>atatatctgacacgtgcctggagactagggagtaatccccttggcggttaaaacgcgggggacagcgcg<br>tacgtgcgtttaagcggtgctagagctgtctacgaccaattgagcggcctcggcaccgggattctccaaa<br>aaaaaaaaaaaactcgagcatcggaggacagtcctccgcggacctgcaggcatgcaagc | Forward:<br>5’GACTGGCACGCTAGCGG<br>Reverse:<br>5’GGACTGTCCTCCGATGC |
| 35-NC-35T$_{15}$ | pGEM-3Z/BNB-601-T15<br>5’gctcggtacccggggatcctatccgactggcacgctagcggtcgccgttcaatagatctaccggatgt<br>atatatctgacacgtgcctggagactagggagtaatccccttggcggttaaaacgcgggggacagcgcg<br>tacgtgcgtttaagcggtgctagagctgtctacgaccaattgagcggcctcggcaccgggattcctttttt<br>ttttttttttctcgagcatcggaggacagtcctccgcggacctgcaggcatgcaagc | Forward:<br>5’GACTGGCACGCTAGCGG<br>Reverse:<br>5’GGACTGTCCTCCGATGC |

| | | |
|---|---|---|
| 35$A_{15}$-NC-35$T_{15}$ | pGEM-3Z/BNB-A15-601-T15<br>5’gctcggtacccggggatcctatccgactggcacgctagcggtcgccaaaaaaaaaaaaaaacggat<br>gtatatatctgacacgtgcctggagactagggagtaatccccttggcggttaaaacgcgggggacagcg<br>cgtacgtgcgtttaagcggtgctagagctgtctacgaccaattgagcggcctcggcaccgggattctcctt<br>tttttttttttttctcgagcatcggaggacagtcctccgcggacctgcaggcatgcaagc | Forward:<br>5’GACTGGCACGCTAGCGG<br>Reverse:<br>5’GGACTGTCCTCCGATGC |
| 35$T_{15}$-NC-35$A_{15}$ | pGEM-3Z/BNB-T15-601-A15<br>5’gctcggtacccggggatcctatccgactggcacgctagcggtcgccttttttttttttttcggatgtatat<br>atctgacacgtgcctggagactagggagtaatccccttggcggttaaaacgcgggggacagcgcgtacg<br>tgcgtttaagcggtgctagagctgtctacgaccaattgagcggcctcggcaccgggattctccaaaaaaa<br>aaaaaaaactcgagcatcggaggacagtcctccgcggacctgcaggcatgcaagc | Forward:<br>5’GACTGGCACGCTAGCGG<br>Reverse:<br>5’GGACTGTCCTCCGATGC |
| 0-NC-140 | pGEM-3Z/601-Gal4:<br>5’ggtcgctgttcaatacatgcacaggatgtatatatctgacacgtgcctggagactagggagtaatccc<br>cttggcggttaaaacgcgggggacagcgcgtacgtgcgtttaagcggtgctagagctgtctacgaccaat<br>tgagcggcctcggcaccgggattctccagggcggccgcgtatctcgagcatcggaggacagtcctccgc<br>ggacctgcaggcatgcaagcttgagtattctatagtgtcacctaaatagcttggcgtaatcatggtcatag<br>ctgtttcctgtgtgaaattgttatccgctcacaattccacacaacatac | Forward:<br>5’ACAGGATGTATATATCTGA<br>CACGTGCCTGG<br>Reverse:<br>5’AGCGGATAACAATTTCACA<br>C |
| 70$A_{15}$-NC-70$T_{15}$ | pGEM-3Z/BNB-A15-601-T15<br>5’tgtaatacgactcactatagggcgaattcgagctcggtacccggggatcctatccgactggcacgcta<br>gcggtcgccaaaaaaaaaaaaaaacggatgtatatatctgacacgtgcctggagactagggagtaatc<br>cccttggcggttaaaacgcgggggacagcgcgtacgtgcgtttaagcggtgctagagctgtctacgacc<br>aattgagcggcctcggcaccgggattctcctttttttttttttttctcgagcatcggaggacagtcctccgcg<br>gacctgcaggcatgcaagcttgagtattctatagtgtcacctaaatagc | Forward:<br>5’GGCGAATTCGAGCTCGGT<br>Reverse:<br>5’GAATACTCAAGCTTGCATG<br>CCTG |
| 70$T_{15}$-NC-70$A_{15}$ | pGEM-3Z/BNB-T15-601-A15<br>5’tgtaatacgactcactatagggcgaattcgagctcggtacccggggatcctatccgactggcacgcta<br>gcggtcgccttttttttttttttcggatgtatatatctgacacgtgcctggagactagggagtaatccccttg<br>gcggttaaaacgcgggggacagcgcgtacgtgcgtttaagcggtgctagagctgtctacgaccaattga<br>gcggcctcggcaccgggattctccaaaaaaaaaaaaaaactcgagcatcggaggacagtcctccgcgg<br>acctgcaggcatgcaagcttgagtattctatagtgtcacctaaatagc | Forward:<br>5’GGCGAATTCGAGCTCGGT<br>Reverse:<br>5’GAATACTCAAGCTTGCATG<br>CCTG |
| 30-NC-70 | pGEM-3Z/BNB-601<br>5’gctcggtacccggggatcctatccgactggcacgctagcggtcgccgttcaatagatctaccggatgt<br>atatatctgacacgtgcctggagactagggagtaatccccttggcggttaaaacgcgggggacagcgcg<br>tacgtgcgtttaagcggtgctagagctgtctacgaccaattgagcggcctcggcaccgggattctccagg<br>gcggccgcgtatctcgagcatcggaggacagtcctccgcggacctgcaggcatgcaagcttgagtattct<br>atagtgtcacctaaatagc | Forward:<br>5’GCACGCTAGCGGTCG<br>Reverse:<br>5’GAATACTCAAGCTTGCATG<br>CCTG |
| 30$A_{15}$-NC-70 | pGEM-3Z/BNB-A15-601<br>5’gctcggtacccggggatcctatccgactggcacgctagcggtcgccaaaaaaaaaaaaaaacggat<br>gtatatatctgacacgtgcctggagactagggagtaatccccttggcggttaaaacgcgggggacagcg<br>cgtacgtgcgtttaagcggtgctagagctgtctacgaccaattgagcggcctcggcaccgggattctcca<br>gggcggccgcgtatctcgagcatcggaggacagtcctccgcggacctgcaggcatgcaagcttgagtat<br>tctatagtgtcacctaaatagc | Forward:<br>5’GCACGCTAGCGGTCG<br>Reverse:<br>5’GAATACTCAAGCTTGCATG<br>CCTG |
| 70-NC-30 | pGEM-3Z/BNB-601<br>5’tgtaatacgactcactatagggcgaattcgagctcggtacccggggatcctatccgactggcacgcta<br>gcggtcgccgttcaatagatctaccggatgtatatatctgacacgtgcctggagactagggagtaatccc<br>cttggcggttaaaacgcgggggacagcgcgtacgtgcgtttaagcggtgctagagctgtctacgaccaat<br>tgagcggcctcggcaccgggattctccagggcggccgcgtatctcgagcatcggaggacagtcctccgc<br>ggacctgcaggcatgcaagc | Forward:<br>5’GGCGAATTCGAGCTCGGT<br>Reverse:<br>5’ GTCCTCCGATGCTCG |
| 70-NC-30$A_{15}$ | pGEM-3Z/BNB-601-A15<br>5’tgtaatacgactcactatagggcgaattcgagctcggtacccggggatcctatccgactggcacgcta<br>gcggtcgccgttcaatagatctaccggatgtatatatctgacacgtgcctggagactagggagtaatccc<br>cttggcggttaaaacgcgggggacagcgcgtacgtgcgtttaagcggtgctagagctgtctacgaccaat<br>tgagcggcctcggcaccgggattctccaaaaaaaaaaaaaaactcgagcatcggaggacagtcctccg<br>cggacctgcaggcatgcaagc | Forward:<br>5’GGCGAATTCGAGCTCGGT<br>Reverse:<br>5’ GTCCTCCGATGCTCG |
| 70-NC-30$T_{15}$ | pGEM-3Z/BNB-601-T15<br>5’tgtaatacgactcactatagggcgaattcgagctcggtacccggggatcctatccgactggcacgcta<br>gcggtcgccgttcaatagatctaccggatgtatatatctgacacgtgcctggagactagggagtaatccc<br>cttggcggttaaaacgcgggggacagcgcgtacgtgcgtttaagcggtgctagagctgtctacgaccaat<br>tgagcggcctcggcaccgggattctcctttttttttttttttctcgagcatcggaggacagtcctccgcggac<br>ctgcaggcatgcaagc | Forward:<br>5’GGCGAATTCGAGCTCGGT<br>Reverse:<br>5’ GTCCTCCGATGCTCG |

| | | |
|---|---|---|
| 5s | pGEX-6P-1/5s<br>5’aggggcccctgggatccccggaattccaacgaataacttccagggatttataagccgatgacgtcata<br>acatccctgaccctttaaatagcttaactttcatcaagcaagagcctacgaccataccatgctgaatatac<br>cggttctcgtccgatcaccgaagtcaagcagcatagggctcggttagtacttggatgggagaccgcctgg<br>gaataccgaattcccgggtcgactcgagcggcc | Forward:<br>5’CAGGGATTTATAAGCCGA<br>TGAC<br>Reverse:<br>5’ACTAACCGAGCCCTATGCT |
| 601 | pGEM-3Z/601-Gal4<br>5’ggtcgctgttcaatacatgcacaggatgtatatatctgacacgtgcctggagactagggagtaatccc<br>cttggcggttaaaacgcgggggacagcgcgtacgtgcgtttaagcggtgctagagctgtctacgaccaat<br>tgagcggcctcggcaccgggattctccagggcggccgcgtatctcgagcatcggaggacagtcctccgc<br>ggacctgcaggcatgcaag | Forward:<br>5’ACAGGATGTATATATCTGA<br>CACGTGCCTGG<br>Reverse:<br>5’CTCGGCACCGGGATTCTCCAG |
| 601A15-SHL1.5 | pGEM-3Z/601-A15-SHL1.5<br>5’ggtcgctgttcaatacatgcacaggatgtatatatctgacacgtgcctggagactagggagtaatccc<br>cttggcggttaaaacgcgggggacagcgcgtacaaaaaaaaaaaaaaagctagagctgtctacgacc<br>aattgagcggcctcggcaccgggattctccagggcggccgcgtatctcgagcatcggaggacagtcctc<br>cgcggacctgcaggcatgcaag | Forward:<br>5’ACAGGATGTATATATCTGA<br>CACGTGCCTGG<br>Reverse:<br>5’CTCGGCACCGGGATTCTCC<br>AG |
| 601T15-SHL1.5 | pGEM-3Z/601-T15-SHL1.5<br>5’ggtcgctgttcaatacatgcacaggatgtatatatctgacacgtgcctggagactagggagtaatccc<br>cttggcggttaaaacgcgggggacagcgcgtactttttttttttttttgctagagctgtctacgaccaattga<br>gcggcctcggcaccgggattctccagggcggccgcgtatctcgagcatcggaggacagtcctccgcgga<br>cctgcaggcatgcaag | Forward:<br>5’ACAGGATGTATATATCTGA<br>CACGTGCCTGG<br>Reverse:<br>5’CTCGGCACCGGGATTCTCCAG |
| 601A15-SHL5.5 | pGEM-3Z/601-A15-SHL5<br>5’ggtcgctgttcaatacatgcacaggatgtatatatctgacacgtgcctggagactagggagtaatccc<br>cttggcggttaaaacgcgggggacagcgcgtacgtgcgtttaagcggtgctagagctgtctacgaccaat<br>tgaaaaaaaaaaaaaaagggattctccagggcggccgcgtatctcgagcatcggaggacagtcctccg<br>cggacctgcaggcatgcaag | Forward:<br>5’ACAGGATGTATATATCTGA<br>CACGTGCCTGG<br>Reverse:<br>5’CTGGAGAATCCCTTTTTTTT |
| 601T15-SHL5.5 | pGEM-3Z/601-T15-SHL5<br>5’ggtcgctgttcaatacatgcacaggatgtatatatctgacacgtgcctggagactagggagtaatccc<br>cttggcggttaaaacgcgggggacagcgcgtacgtgcgtttaagcggtgctagagctgtctacgaccaat<br>tgtttttttttttttttgggattctccagggcggccgcgtatctcgagcatcggaggacagtcctccgcggac<br>ctgcaggcatgcaag | Forward:<br>5’ACAGGATGTATATATCTGA<br>CACGTGCCTGG<br>Reverse:<br>5’CTGGAGAATCCCAAAAAAAA |
| 0-NC-30 | pGEM-3Z/601-Gal4<br>5’ggtcgctgttcaatacatgcacaggatgtatatatctgacacgtgcctggagactagggagtaatccc<br>cttggcggttaaaacgcgggggacagcgcgtacgtgcgtttaagcggtgctagagctgtctacgaccaat<br>tgagcggcctcggcaccgggattctccagggcggccgcgtatctcgagcatcggaggacagtcctccgc<br>ggacctgcaggcatgcaag | Forward:<br>5’ACAGGATGTATATATCTGA<br>CACGTGCCTGG<br>Reverse:<br>5’GTCCTCCGATGCTCG |
| 0-NC-30A15 | pGEM-3Z/601-Gal4-A15-2in<br>5’ggtcgctgttcaatacatgcacaggatgtatatatctgacacgtgcctggagactagggagtaatccc<br>cttggcggttaaaacgcgggggacagcgcgtacgtgcgtttaagcggtgctagagctgtctacgaccaat<br>tgagcggcctcggcaccgggattctccaaaaaaaaaaaaaaactcgagcatcggaggacagtcctccg<br>cggacctgcaggcatgcaag | Forward<br>5’ACAGGATGTATATATCTGA<br>CACGTGCCTGG<br>Reverse<br>5’GTCCTCCGATGCTCG |
| 0-NC-30T15 | pGEM-3Z/601-Gal4-T15-2in<br>5’ggtcgctgttcaatacatgcacaggatgtatatatctgacacgtgcctggagactagggagtaatccc<br>cttggcggttaaaacgcgggggacagcgcgtacgtgcgtttaagcggtgctagagctgtctacgaccaat<br>tgagcggcctcggcaccgggattctcctttttttttttttttctcgagcatcggaggacagtcctccgcggac<br>ctgcaggcatgcaag | Forward<br>5’ACAGGATGTATATATCTGA<br>CACGTGCCTGG<br>Reverse<br>5’GTCCTCCGATGCTCG |

Letters in red correspond to the nucleosome positioning region of Widom’s 601 sequence (1).

Letters in blue correspond to the dominant positioning region of the *Lytechinus variegatus* 5s rDNA (2, 3).

Sequences highlighted in yellow correspond to the regions bound by the PCR primers.

Regions highlighted in cyan correspond to flanking sequences in the template plasmid that match - and are continued by - the original Widom’s pGEM-3Z/601 plasmid sequence (4).

Regions highlighted in green correspond to flanking sequences that match -and are continued by- the pGEX-6P-1 plasmid sequence.


#### Table S2. Yeast strains used for purification of ATP-dependent chromatin remodeling complexes.

| Complex (tagged subunit) | Strain | Name | Genotype (background) | Source |
|---|---|---|---|---|
| **RSC (Rsc2)** | YLR357w | Rsc2-TAP | *RSC2-TAP (BY4741)* | Open Biosystems |
| **ISW1a (Ioc3)** | YFR013w | Ioc3-TAP | *IOC3-TAP (BY4741)* | Open Biosystems |

### Figure S1

**A**

5'--NNN**ACAG**GATGTATATATCTGACACGTGCCTGGAGACTAGGGAGTAATCCCCTTGGCGGTTAAAACGCGGGGGACAGCGCGTACGTGCGTTTAAGCGGTGCTAGAGCTGTCTACGACCAATTGAGCGGCCTCGGCACCGGGATTCTCCAGNNN--3'

3'--NNNTGTCCTACATATATAGACTGTGCACGGACCTCTGATCCCTCATTAGGGGAACCGCCAATTTTGCGCCCCCTGTCGCGCATGCACGCAAATTCGCCACGATCTCGACAGATGCTGGTTAACTCGCCGGAGCCGTGGCCCTAAGAGGTCNNN--5'

601 nucleosome positioning sequence

**B**

Core/dyad located 3' of polyA
(Graphs' bars in green color)

Core/dyad located 5' of polyA
(Graphs' bars in orange color)

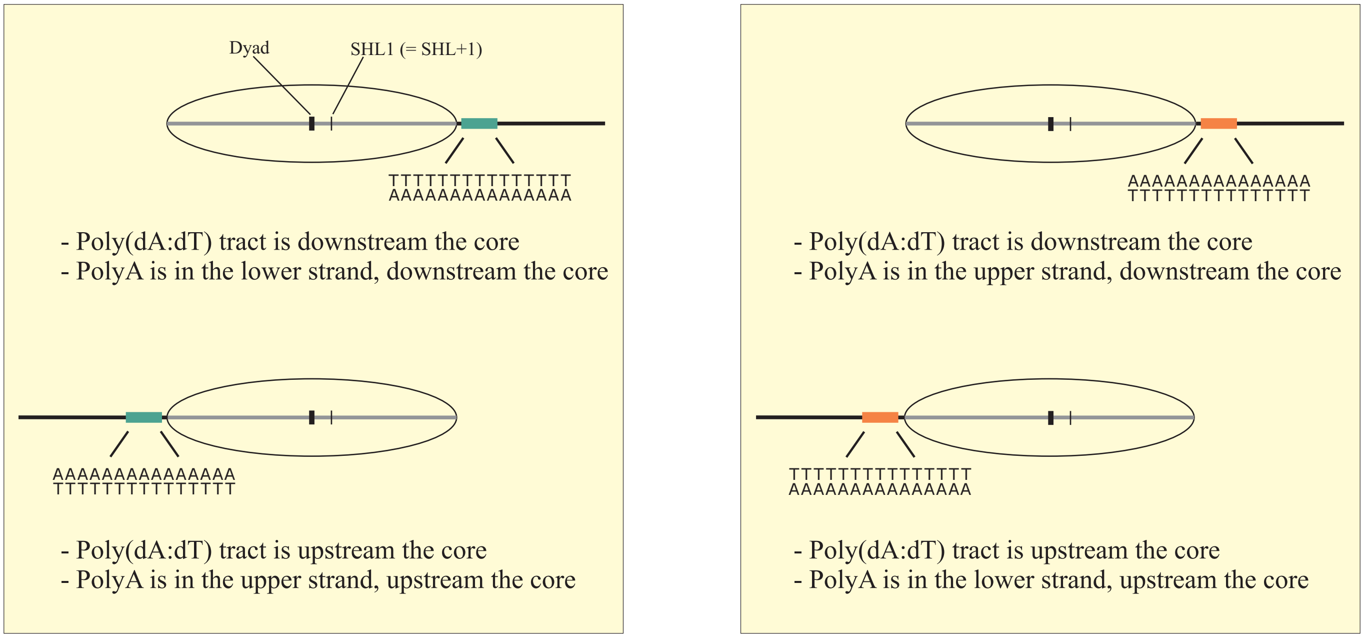


Configuration more frequently found at gene promoters in *S. cerevisiae* for -1 (top) and +1 nucleosomes (bottom), when making the upper strand equivalent to the sense strand of genes

Configuration less frequently found at gene promoters in *S. cerevisiae*, when making the upper strand equivalent to the sense strand of genes

### Figure S2

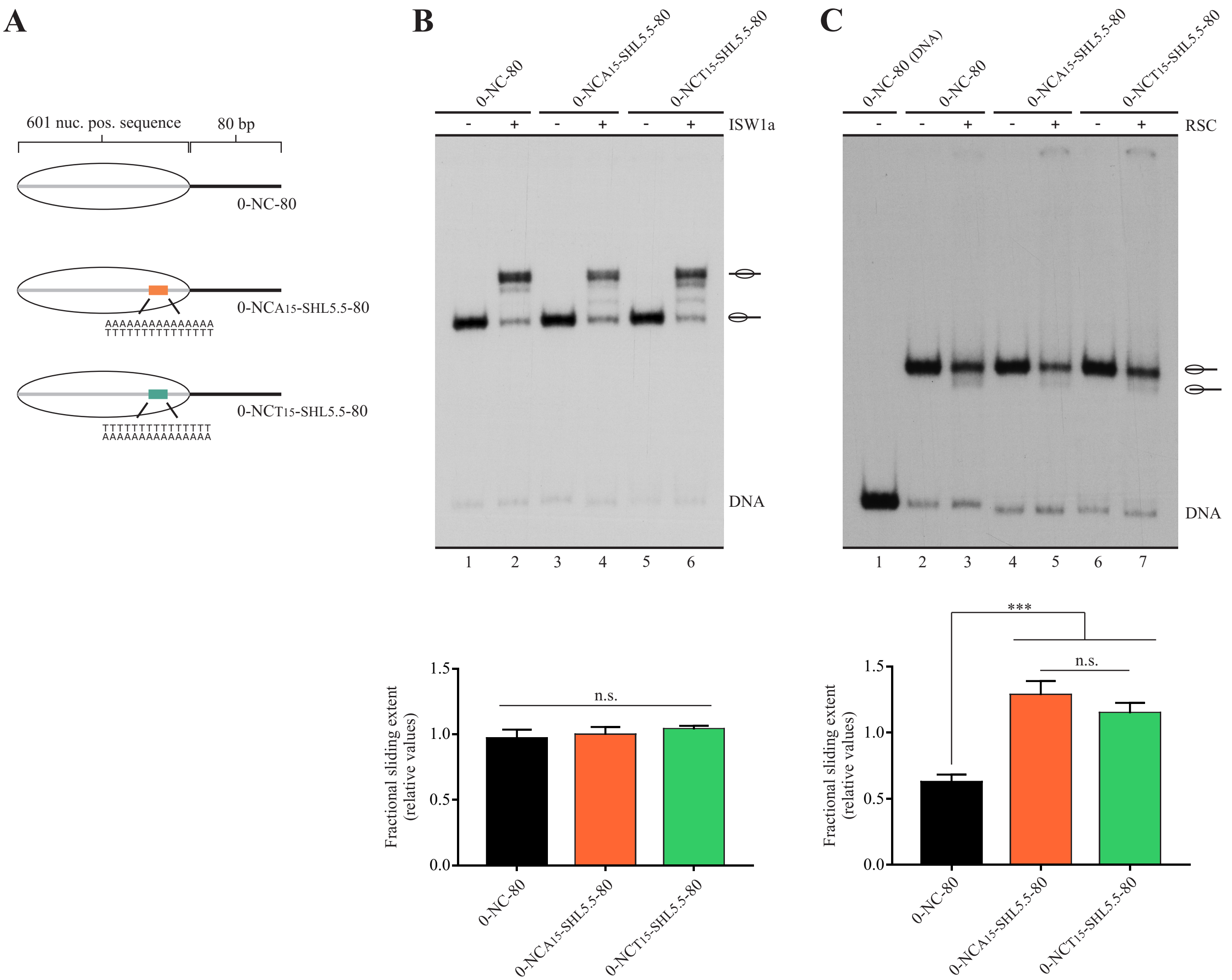
A
601 nuc. pos. sequence
80 bp
0-NC-80
0-NCA15-SHL5.5-80
AAAAAAAAAAAAAAA
TTTTTTTTTTTTTTT
0-NCT15-SHL5.5-80
TTTTTTTTTTTTTTT
AAAAAAAAAAAAAAA
B
0-NC-80
0-NCA15-SHL5.5-80
0-NCT15-SHL5.5-80
ISW1a
DNA
1 2 3 4 5 6
C
0-NC-80 (DNA)
0-NC-80
0-NCA15-SHL5.5-80
0-NCT15-SHL5.5-80
RSC
DNA
1 2 3 4 5 6 7
Fractional sliding extent (relative values)
n.s.
***
1.5
1.0
0.5
0.0

### Figure S3

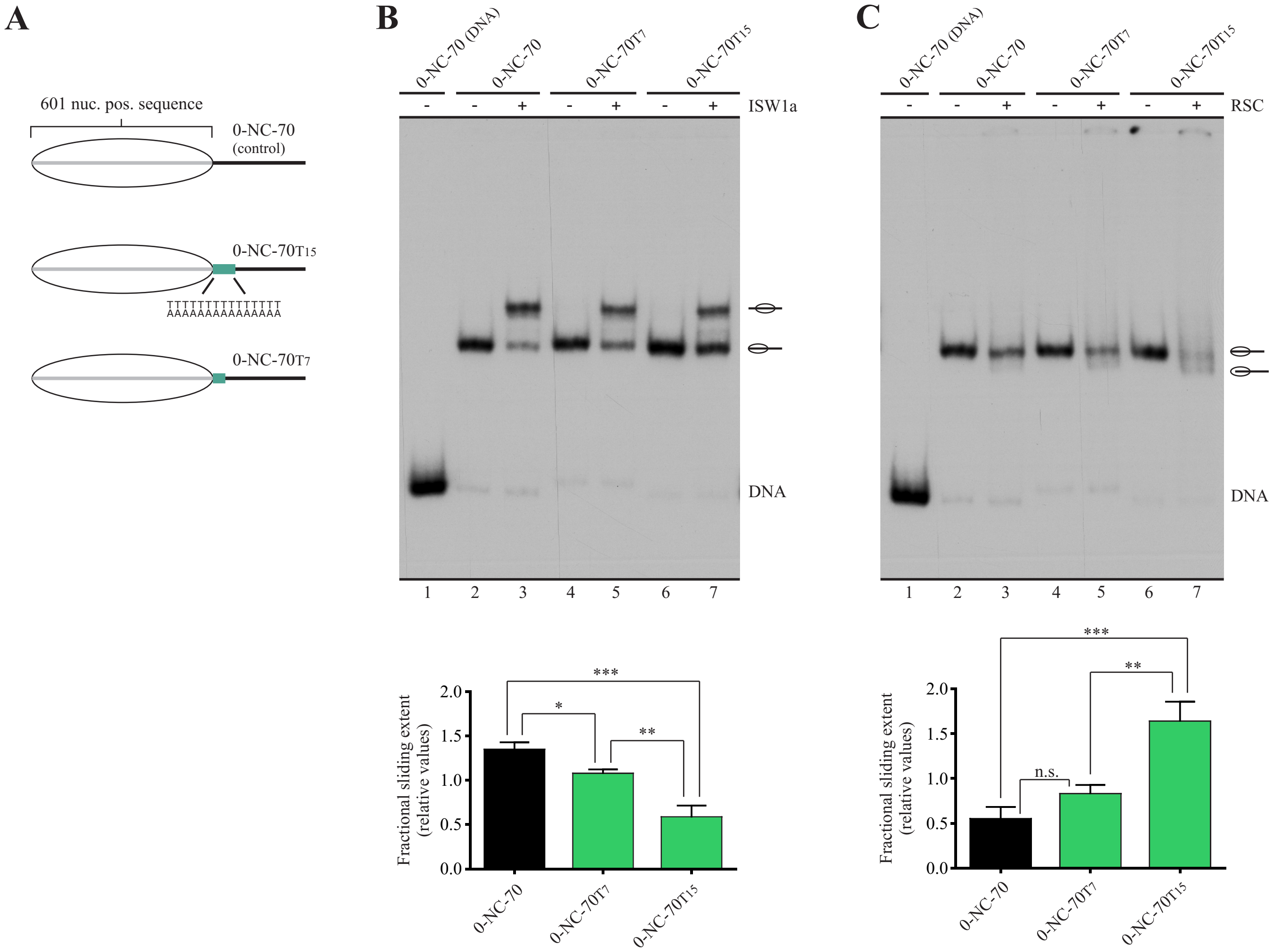

A
B
C
601 nuc. pos. sequence
0-NC-70
(control)
0-NC-70T15
TTTTTTTTTTTTTTT
AAAAAAAAAAAAAAA
0-NC-70T7
0-NC-70 (DNA)
0-NC-70
0-NC-70T7
0-NC-70T15
ISW1a
RSC
DNA
1 2 3 4 5 6 7
Fractional sliding extent
(relative values)
0.0
0.5
1.0
1.5
2.0
*
**
***
n.s.

### Figure S4

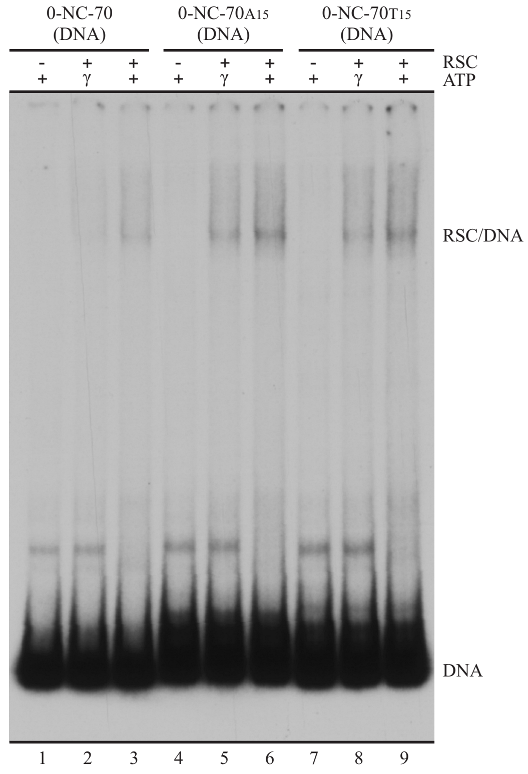

0-NC-70 (DNA)
0-NC-70A15 (DNA)
0-NC-70T15 (DNA)
RSC
ATP
- + + - + + - + +
+ γ + + γ + + γ +
RSC/DNA
DNA
1 2 3 4 5 6 7 8 9

### Figure S5

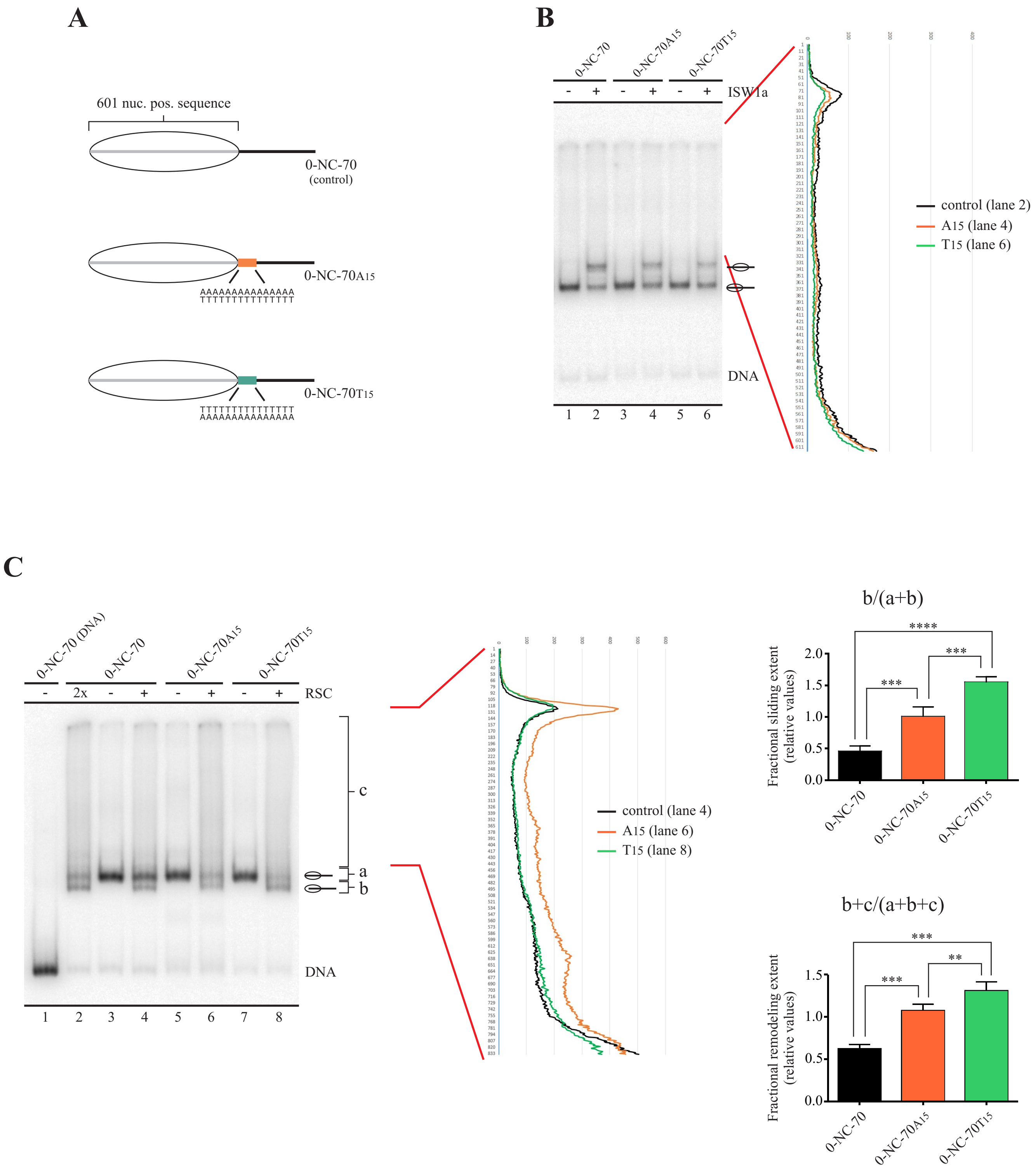
A
601 nuc. pos. sequence
0-NC-70
(control)
0-NC-70A15
AAAAAAAAAAAAAAA
TTTTTTTTTTTTTTT
0-NC-70T15
TTTTTTTTTTTTTTT
AAAAAAAAAAAAAAA
B
0-NC-70
0-NC-70A15
0-NC-70T15
- + - + - + ISW1a
DNA
1 2 3 4 5 6
control (lane 2)
A15 (lane 4)
T15 (lane 6)
C
0-NC-70 (DNA)
0-NC-70
0-NC-70A15
0-NC-70T15
- 2x - + - + - + RSC
c
a
b
DNA
1 2 3 4 5 6 7 8
control (lane 4)
A15 (lane 6)
T15 (lane 8)
b/(a+b)
Fractional sliding extent
(relative values)
0-NC-70
0-NC-70A15
0-NC-70T15
b+c/(a+b+c)
Fractional remodeling extent
(relative values)
0-NC-70
0-NC-70A15
0-NC-70T15

### Figure S6

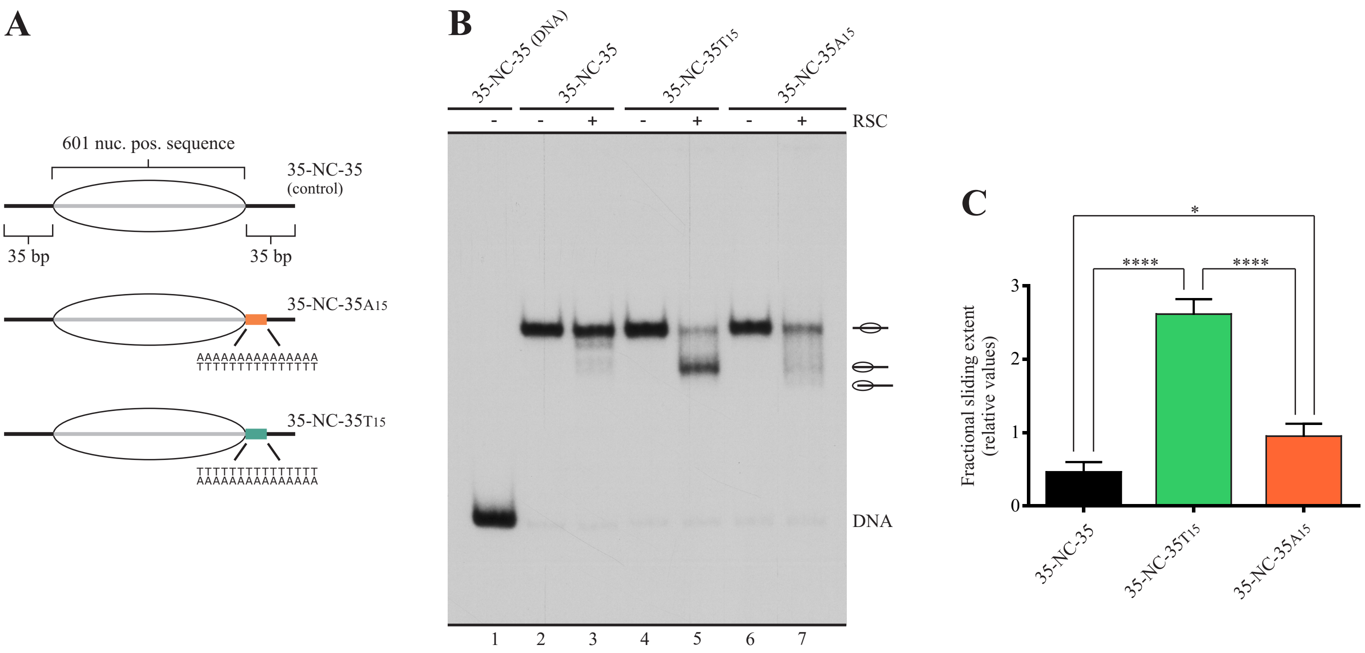

A
601 nuc. pos. sequence
35-NC-35
(control)
35 bp
35 bp
35-NC-35A15
AAAAAAAAAAAAAAA
TTTTTTTTTTTTTTT
35-NC-35T15
TTTTTTTTTTTTTTT
AAAAAAAAAAAAAAA
B
35-NC-35 (DNA)
35-NC-35
35-NC-35T15
35-NC-35A15
RSC
- - + - + - +
DNA
1 2 3 4 5 6 7
C
Fractional sliding extent
(relative values)
*
****
****
0
1
2
3
35-NC-35
35-NC-35T15
35-NC-35A15

### Figure S7

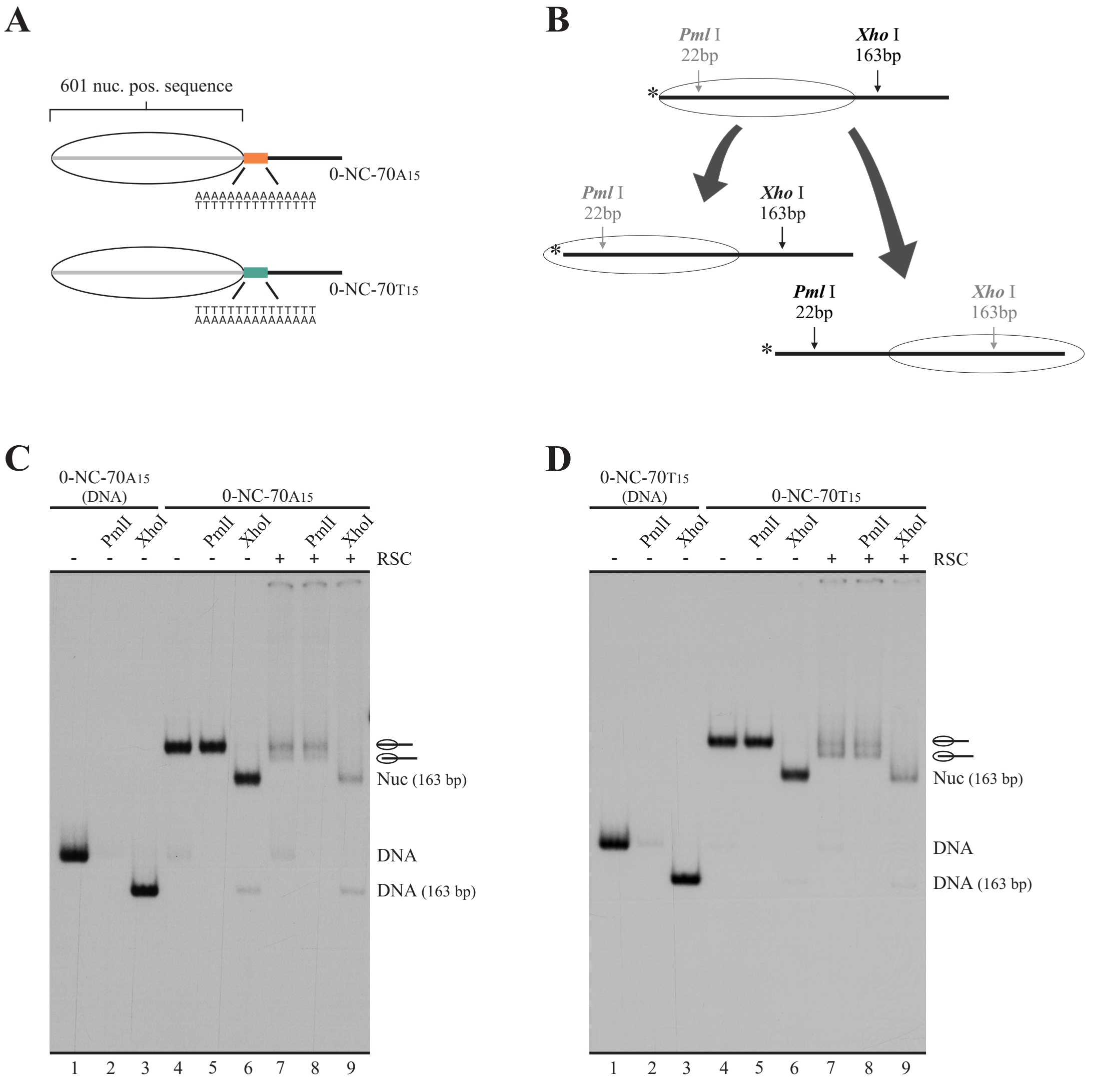
A
601 nuc. pos. sequence
AAAAAAAAAAAAAAA
TTTTTTTTTTTTTTT
0-NC-70A15
0-NC-70T15
B
Pml I
22bp
Xho I
163bp
C
0-NC-70A15
(DNA)
0-NC-70A15
PmlI
XhoI
RSC
Nuc (163 bp)
DNA
DNA (163 bp)
1 2 3 4 5 6 7 8 9
D
0-NC-70T15
(DNA)
0-NC-70T15
PmlI
XhoI
RSC
Nuc (163 bp)
DNA
DNA (163 bp)
1 2 3 4 5 6 7 8 9

### Figure S8

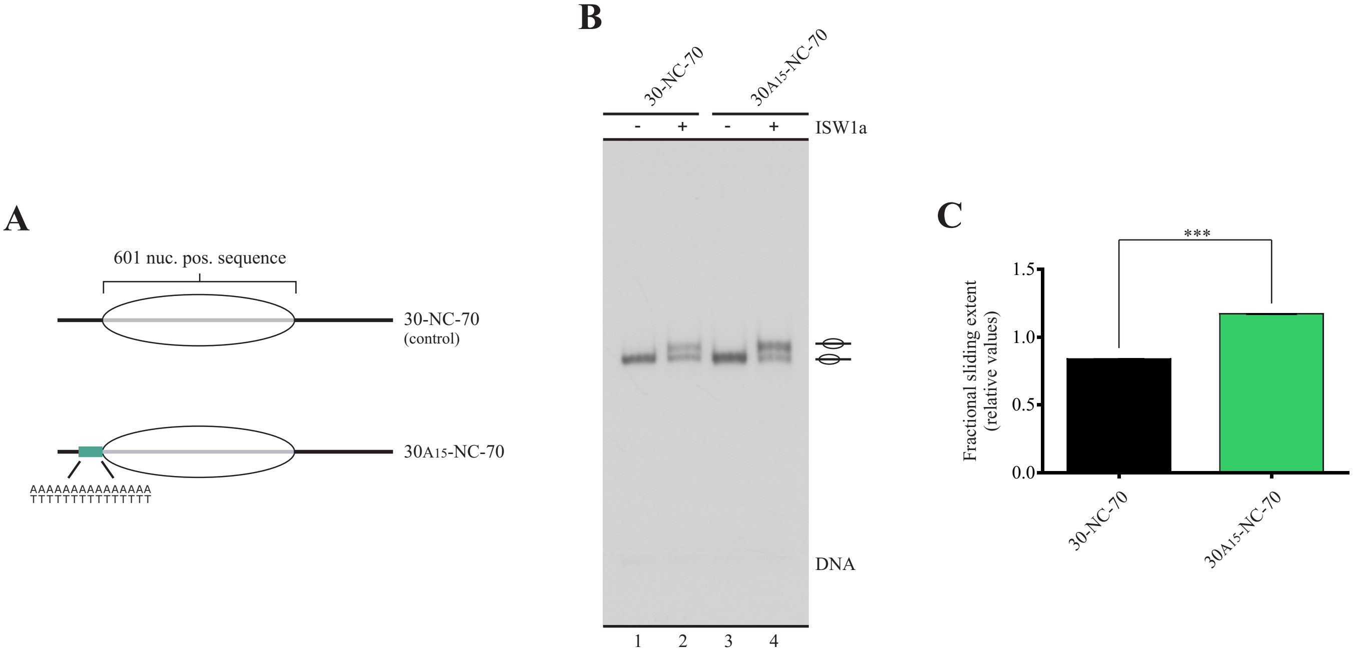
A
601 nuc. pos. sequence
30-NC-70
(control)
30A15-NC-70
AAAAAAAAAAAAAAA
TTTTTTTTTTTTTTT
B
30-NC-70
30A15-NC-70
- + - +
ISW1a
DNA
1 2 3 4
C
***
Fractional sliding extent
(relative values)
1.5
1.0
0.5
0.0
30-NC-70
30A15-NC-70

### Figure S9

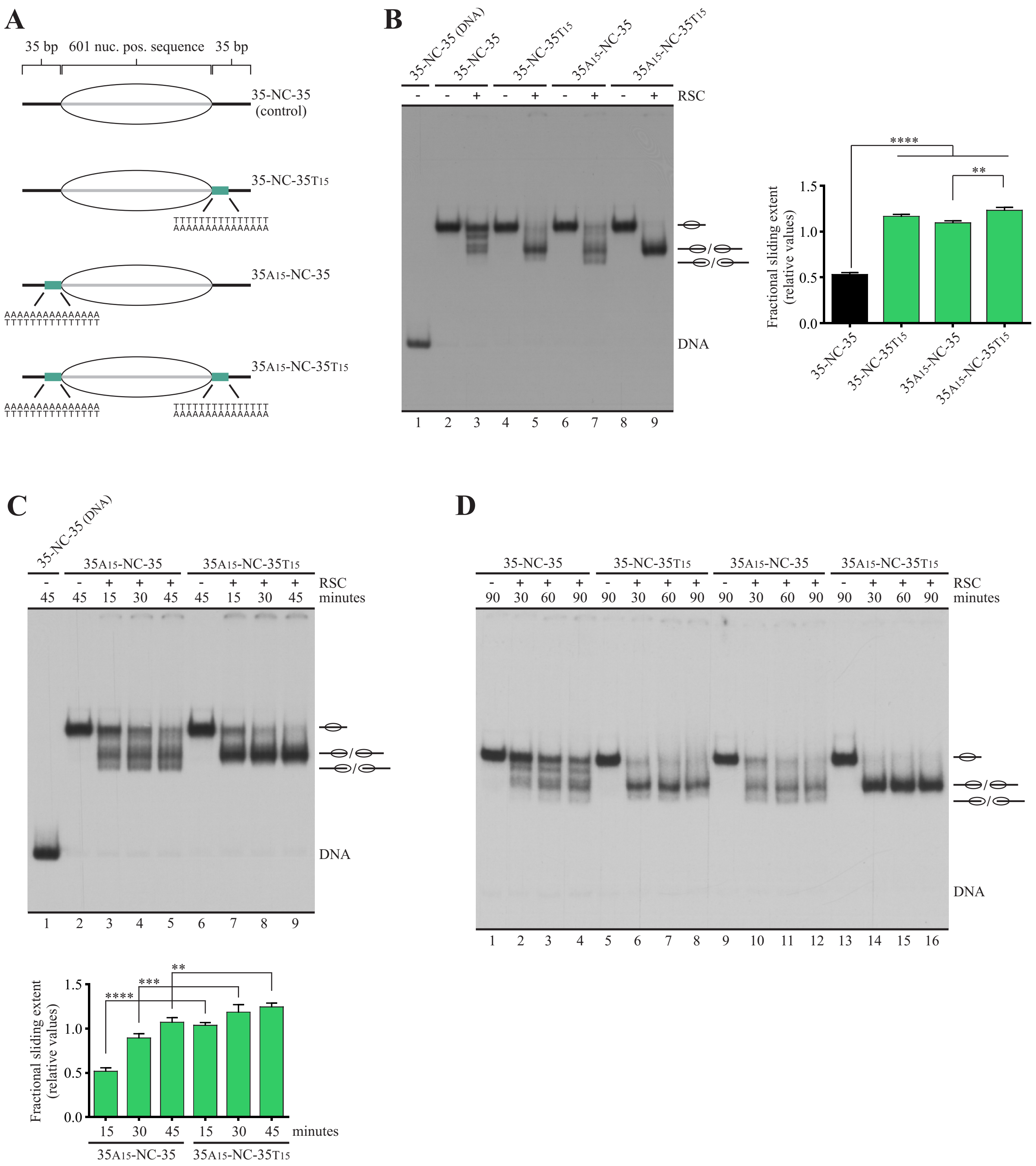

A
35 bp
601 nuc. pos. sequence
35 bp
35-NC-35
(control)
35-NC-35T15
35A15-NC-35
35A15-NC-35T15
B
35-NC-35 (DNA)
35-NC-35
35-NC-35T15
35A15-NC-35
35A15-NC-35T15
RSC
DNA
1 2 3 4 5 6 7 8 9
Fractional sliding extent
(relative values)
1.5
1.0
0.5
0.0
****
**
C
35-NC-35 (DNA)
35A15-NC-35
35A15-NC-35T15
RSC
minutes
45 45 15 30 45 45 15 30 45
DNA
1 2 3 4 5 6 7 8 9
****
***
**
15 30 45 15 30 45 minutes
D
35-NC-35
35-NC-35T15
35A15-NC-35
35A15-NC-35T15
RSC
minutes
90 30 60 90 90 30 60 90 90 30 60 90 90 30 60 90
DNA
1 2 3 4 5 6 7 8 9 10 11 12 13 14 15 16
